# Collagen type VI regulates TGFβ bioavailability in skeletal muscle

**DOI:** 10.1101/2023.06.22.545964

**Authors:** Payam Mohassel, Jachinta Rooney, Yaqun Zou, Kory Johnson, Gina Norato, Hailey Hearn, Matthew A Nalls, Pomi Yun, Tracy Ogata, Sarah Silverstein, David A Sleboda, Thomas J Roberts, Daniel B Rifkin, Carsten G Bönnemann

## Abstract

Collagen VI-related disorders (*COL6*-RDs) are a group of rare muscular dystrophies caused by pathogenic variants in collagen VI genes (*COL6A1, COL6A2,* and *COL6A3*). Collagen type VI is a heterotrimeric, microfibrillar component of the muscle extracellular matrix (ECM), predominantly secreted by resident fibroadipogenic precursor cells in skeletal muscle. The absence or mislocalizatoion of collagen VI in the ECM underlies the non-cell autonomous dysfunction and dystrophic changes in skeletal muscle with an as of yet elusive direct mechanistic link between the ECM and myofiber dysfunction. Here, we conduct a comprehensive natural history and outcome study in a novel mouse model of *COL6*-RDs (*Col6a2^-/-^*mice) using standardized (Treat-NMD) functional, histological, and physiologic parameter. Notably, we identify a conspicuous dysregulation of the TGFβ pathway early in the disease process and propose that the collagen VI deficient matrix is not capable of regulating the dynamic TGFβ bioavailability at baseline and also in response to muscle injury. Thus, we propose a new mechanism for pathogenesis of the disease that links the ECM regulation of TGFβ with downstream skeletal muscle abnormalities, paving the way for developing and validating therapeutics that target this pathway.

## Introduction

Collagen VI-related disorders (*COL6*-RDs) are a group of muscular dystrophies that combine clinical features of connective tissue diseases and muscular dystrophy^1^ and clinically manifest with progressive skeletal muscle weakness and fibrosis, joint contractures, skin abnormalities, osteopenia, and respiratory insufficiency^2, 3^. While progress has been made in understanding the underlying molecular mechanisms of *COL6-*RDs, no approved disease modifying therapies are available and treatment relies on symptomatic management.

It is notable that mild and non-progressive muscle weakness is a common finding in genetic connective tissue diseases (for example, Marfan’s syndrome due to *FBN1* pathogenic variants^4^, osteogenesis imperfecta due to *COL1A1* pathogenic variants^5^, and Ehlers-Danlos/myopathy overlap due to *COL12A1* pathogenic variants^6^). However, *COL6-*RDs are unique as they present not only with muscle weakness, but also with a progressive, degenerative disease characterized by dystrophic changes and muscle fibrosis.

Collagen type VI is a microfibrillar component of the extracellular matrix (ECM), with suprastructural similarities to fibrillin microfibrils with which it often co-purifies^7, 8^. Its expression is widespread in brain, blood vessels, skin, cardiac and skeletal muscle, connective tissues (for example, cartilage, bone, tendon, ligaments, and joints), as well as cornea and placenta^1^. Collagen VI is assembled intracellularly from three alpha chains encoded by *COL6A1*, *COL6A2*, and *COL6A3*. Three other genes (*COL6A4—*a pseudogene in humans, *COL6A5*, and *COL6A6*) share homology with *COL6A3*^9, 10^, though pathogenic variants in these genes leading to *COL6-*RD have yet to be identified and thus their relevance to this disease remains unclear. The collagen VI ɑ1, ɑ2, and ɑ3 chains each contain a short triple helical domain and assemble from the C-terminal to N-terminal direction into a heterotrimeric monomer. Heterotrimeric monomers form anti- parallel dimers, which subsequently assemble into tetramers before they are secreted into the extracellular space where they join to form a microfibrillar structure in the skeletal muscle basement membrane. Variants that cause the absence of any of the essential ⍺-chains (typically biallelic) or result in assembly-competent mutant chains (typically monoallelic) most often underlie the disease^11^.

How abnormalities in collagen VI in the muscle ECM lead to muscular dystrophy remain incompletely understood. Sarcolemmal fragility and impaired membrane repair, abnormalities in muscle enzymes, intermediate filaments or nuclear envelope proteins, or myofibrillar or Z-disk associated proteins underlie different muscular dystrophies^12, 13^. However, in *COL6-*RDs, these mechanisms have not been shown to be the primary drivers of disease and serum creatine kinase levels are consistently normal or only mildly elevated in *COL6-*RDs^14^. In addition, collagen VI is predominantly produced and secreted by skeletal muscle interstitial fibroblasts and fibroadipogenic precursor cells and not muscle fibers themselves^15^, which further adds to the complexities of *COL6-*RD pathogenesis.

A few mouse models of *COL6*-RDs have been generated and characterized so far: *Col6a1* deficient mice^16, 17^, *Col6a3* deficient mice^18^, and two dominant negative mutant mouse models with deletion of exon 16 of *Col6a3* or exon 5 of *Col6a2*^19, 20^. These mouse models show variable myopathic changes and abnormalities, albeit milder than those of individuals with analogous pathogenic variants and *COL6-*RD. In the most extensively studied mouse model, *Col6a1* knockout mice generated by Bonaldo et al^16^, a few downstream cell physiological disturbances in skeletal muscle have been reported. These include reduction of autophagy flux^21^, mitochondrial abnormalities and apoptosis^22^, and reduced satellite cell regenerative reserve^23, 24^. However, the underlying mechanism of how collagen VI abnormalities in the muscle extracellular matrix leads to these downstream cellular abnormalities and muscular dystrophy has not been fully delineated.

To add to the models of the disease and better understand the underlying mechanisms of *COL6-*RD, we derived and characterized a novel mouse model, *Col6a2^-/-^* mice, from the UC Davis Knockout Mouse project. To establish a robust preclinical model for mechanistic and therapeutic studies, we have comprehensively characterized the natural history of disease in this mouse model using behavioral, physiologic, and histologic studies of skeletal muscle. We identify a conspicuous dysregulation of the TGFβ pathway early in the disease process and propose that the collagen VI deficient matrix is not capable of providing dynamic TGFβ bioavailability and thus propose a new mechanism for pathogenesis of the disease that links the extracellular matrix with downstream skeletal muscle abnormalities.

## Results

### *Col6a2^-/-^* mice have early post-natal muscle atrophy and muscle weakness

We obtained and backcrossed a previously uncharacterized mouse strain with complete deletion of *Col6a2* genomic locus from the UC Davis knockout mouse project repository (Figure 1A). Unlike wildtype and heterozygous (*Col6a2^+/-^*) littermates, mice with homozygous deletion of *Col6a2* (*Col6a2^-/-^*) had complete absence of collagen VI staining in skeletal muscle ECM (Figure 1B). *Col6a2^-/-^* mice weighed less and appeared smaller than littermates (Figure 1C), first noted at around four weeks of age. They had a normal life span and were present in slightly reduced Mendelian ratios (20.6%) (Supplementary Figure 1). Isolated *Col6a2^-/-^*mouse skeletal muscles weighed less than those of littermates and remained proportionally atrophic in mice as old as 60 weeks (Figure 1D).

**Figure 1.**
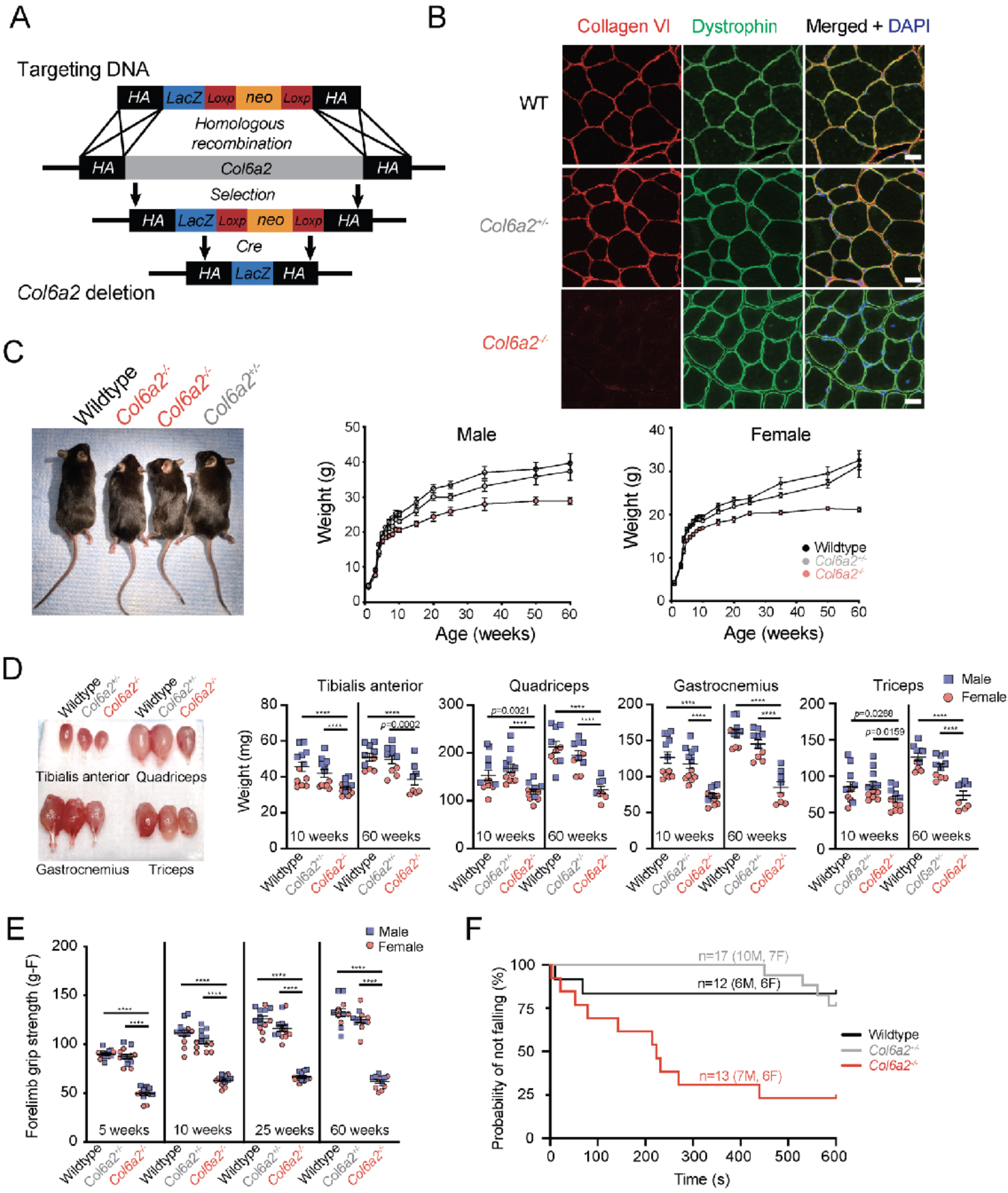
*Col6a2^-/-^* mice develop early post-natal muscle atrophy and weakness. **(A)** The schematic representation of the *Col6a2* knockout allele. The *LacZ* sequence is not expressed but is used in the genotyping strategy. **(B)** Immunofluorescence staining of frozen gastrocnemius muscle shows complete absence of collagen VI from the muscle extracellular matrix in *Col6a2^-/-^* mice. Scale bar= 25 µm **(C)** *Col6a2^-/-^* mice are smaller and weigh less than wildtype and *Col6a2^+/-^* littermates. **(D)** Male and female *Col6a2^-/-^* mice have smaller muscles compared to wildtype and *Col6a2^+/-^* littermates. **(E)** Male and female *Col6a2^-/-^* mice have markedly reduced forelimb grip strength compared to wildtype and *Col6a2^+/-^* littermates. (**F)** *Col6a2^-/-^* mice, assessed at 10 weeks of age, have an increased probability of falling in the hanging wire test. Statistical comparisons in F were performed using Cox proportional hazard model. Genotype is significantly related to the hazard of falling (log-rank test = 17.9 (df=2); p<0.001), with the following pairwise differences: wildtype vs *Col6a2^-/-^*, HR= 0.12 (95% CI: 0.03-0.57), p=0.02; *Col6a2^+/-^* vs *Col6a2^-/-^*, HR = 0.15 (95% CI: 0.05-0.50), p=0.006; *Col6a2^+/-^* vs wildtype, HR = 1.25 (95% CI: 0.23-6.84), p=1.0. M= male, F= female. For all other panels, error bars indicate SEM and statistical comparisons were performed by 2-way ANOVA with Tukey’s adjustment for multiple comparisons. **** *p<*0.0001

*Col6a2^-/-^* mice had markedly reduced forelimb grip strength as early as five weeks of age through 60 weeks of age compared to controls (Figure 1E) and were more likely to fall in the hanging wire test assessed at 10 weeks of age (Figure 1F). Other functional tests such as rotarod, treadmill running, and activity monitoring were highly variable and failed to identify a difference between *Col6a2^-/-^* mice and littermate controls (Supplementary Figure 1).

### *Col6a2^-/-^* skeletal muscles show mild dystrophic features

Histological analysis of *Col6a2^-/-^* mouse muscle showed dystrophic features including fiber size variability with atrophic and hypertrophic myofibers, increased endomysial fibrosis and an abundance of central nuclei, an indirect marker of myofiber regeneration (Figure 2A). Similar findings including rare degenerating and regenerating myofibers were also observed in gastrocnemius, quadriceps, triceps, and diaphragm muscles (Supplementary Figure 2). Immunostaining for specific myosin subtypes failed to identify selective loss or atrophy of oxidative or glycolytic myofibers (Supplementary Figure 2). There was a substantial increase in internal nuclei in *Col6a2^-/-^* mouse muscle (Figure 2B). Minimal feret diameter of myofibers in gastrocnemius muscle sections were quantitatively assessed using myosoft, an open-source software^25^. In 10-week-old animals, fiber size variation, measured by coefficient of variability (standard deviation of minimal feret diameter x 1000/mean of minimal feret diameter), was markedly increased in *Col6a2^-/-^* muscle compared to heterozygous and wildtype controls (Figure 2C). Similar findings were observed in 60-week-old animals (Supplementary Figure 2).

**Figure 2.**
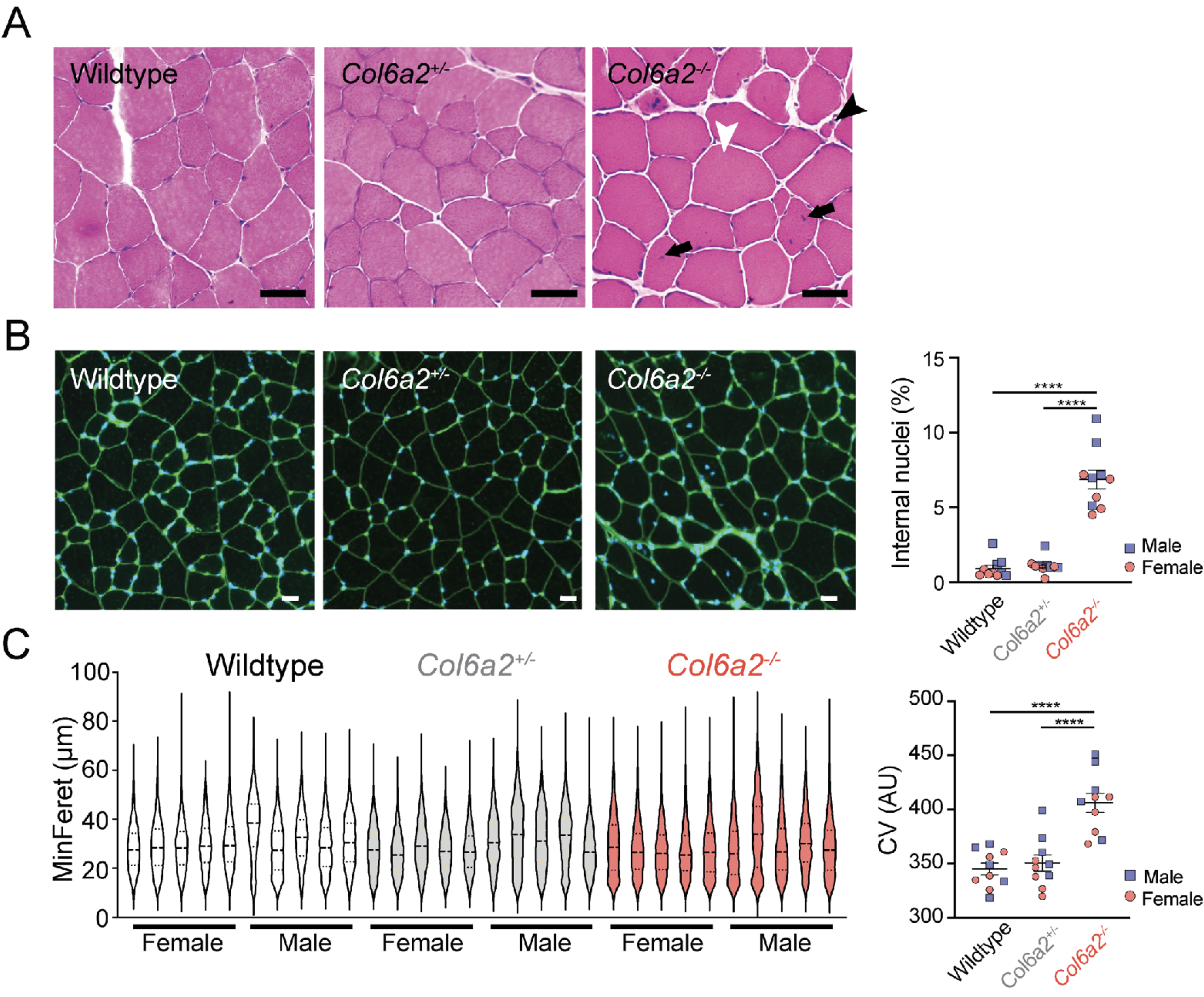
*Col6a2^-/-^* muscle shows myofiber atrophy and dystrophic changes. **(A)** Frozen sections of tibialis anterior muscle of 10-week-old animals were stained with hematoxylin and eosin (H&E). Dystrophic features include fiber size variability with myofiber atrophy (black arrowhead), myofiber hypertrophy (white arrowhead), and increased internal nuclei (arrows). Scale bar = 50 µm. **(B)** Representative images of wheat germ agglutinin (WGA--green) and DAPI stained whole gastrocnemius muscle from 10-week-old male and female mice. Percentage of fibers with internal nuclei were markedly increased in *Col6a2^-/-^* muscle. Scale bar = 50 µm. **(C)** The myofiber minferet diameter of the entire gastrocnemius muscle sections of 10 week old mice were quantified and shown in violin plots. Note reduced mean fiber diameter in *Col6a2^-/-^* mouse muscle and a marked increase in coefficient of variability (CV) consistent with increased number of atrophic and hypertrophic fibers in *Col6a2^-/-^*mouse muscle. AU= arbitrary units. Error bars indicated SEM. Statistical comparisons were performed by 2- way ANOVA and Tukey’s adjustment for multiple comparisons. **** = *p<*0.0001.

### *Col6a2^-/-^* skeletal muscle produces reduced contractile force and is susceptible to lengthening contraction injury

Physiological parameters in a standard *ex vivo* preparation of isolated EDL muscle in 10 week old animals showed a decrease in twitch and maximum tetanic force in *Col6a2^-/-^*muscle (Figure 3A-C). However, specific force, i.e., tetanic force normalized to functional cross-sectional area of the muscle, was only minimally reduced (Figure 3D). We also performed repeated eccentric contractions (10% of optimal length, *L_o_*), which resulted in a marked reduction of the tetanic force in *Col6a2^-/-^* muscle, suggestive of its susceptibility to eccentric contraction injury (Figure 3E). Notably, female *Col6a2^-/-^* muscle showed a less dramatic reduction in tetanic force after repeated eccentric contractions compared to male animals (Figure 3F). Similar contractile force and physiologic parameters were noted in 60-week-old animals and 60-week-old female *Col6a2^-/-^* mice were also protected against tetanic force reduction after repeated eccentric contractions (Supplementary Figure 3). To investigate the susceptibility to lengthening contraction injury *in vivo*, we used a downhill running paradigm followed by Evan’s blue dye (EBD) injection. Consistent with the *ex vivo* physiology testing, we found an increase in sarcolemmal fragility and EBD uptake in *Col6a2^-/-^* muscle fibers after downhill running (Figure 3G-H), though without an apparent protection in female animals.

**Figure 3.**
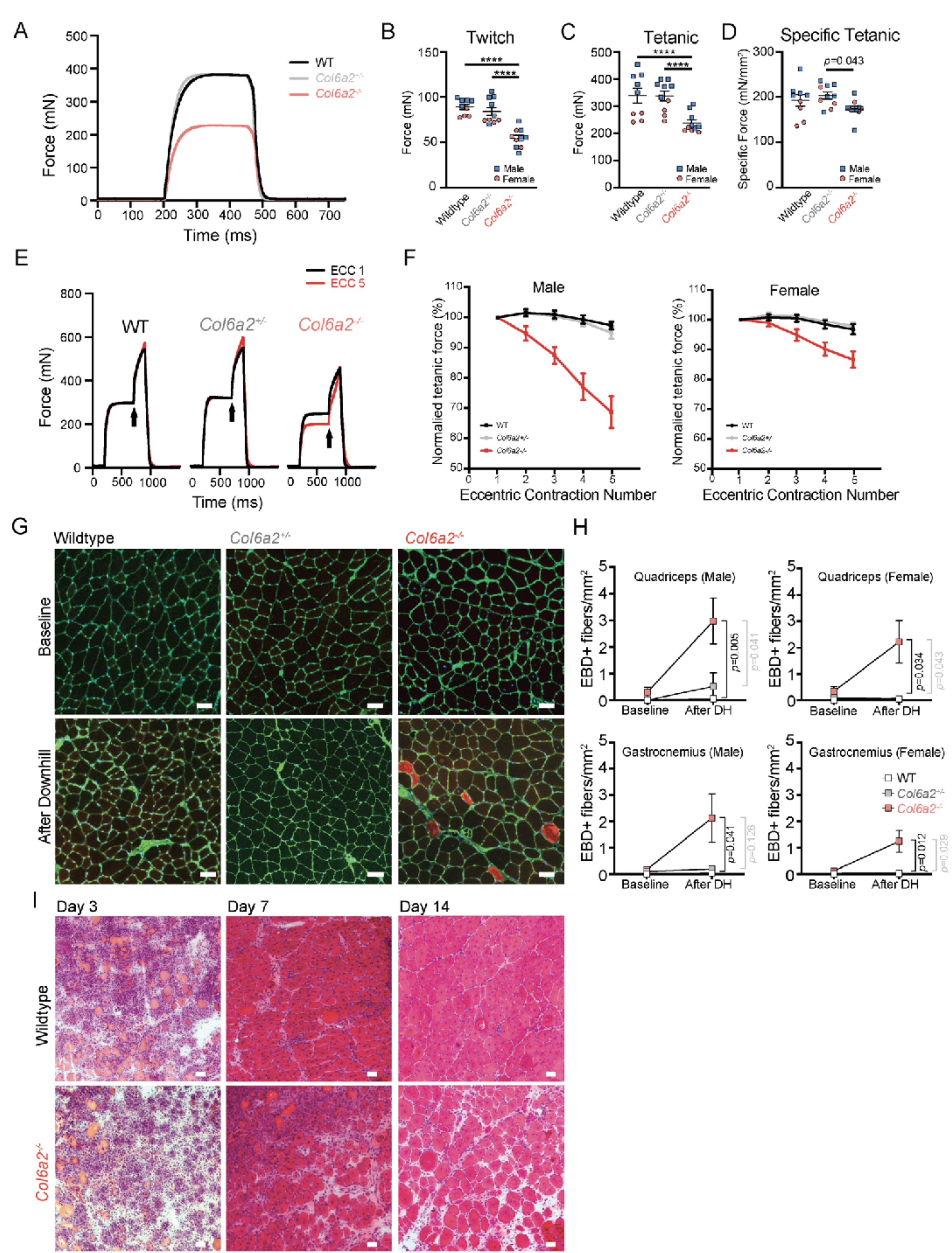
*Col6a2^-/-^*muscle has reduced contractile force, is susceptible to lengthening contraction injury, and shows impaired regenerative capacity. **(A)** Representative tracing of total tetanic force measurement in isolated EDL muscle. (**B)** Maximal twitch force and total tetanic force **(C)** are markedly reduced in 10-week-old *Col6a2^-/-^* mouse EDL muscle. **(D)** The difference in force generation was negligible after normalization to functional cross-sectional area (i.e. specific force). B-D: Error bars indicated SEM. Statistical comparisons were performed by 2-way ANOVA and Tukey’s adjustment for multiple comparisons. **** = *p<*0.0001. **(E)** Representative tracing of force measurement in isolated EDL muscle after eccentric contraction. (F) Tetanic force declined precipitously after repeated eccentric contractions in *Col6a2^-/-^* EDL muscle. The decline is more prominent in male animals. Statistical analysis was performed using linear mixed models and Bonferroni adjustment for multiple comparisons. Male: wildtype vs *Col6a2^--^* (*p<*0.0001); *Col6a2^+/-^*vs *Col6a2^-/-^* (*p<*0.0001). Female: wildtype vs *Col6a2^-/-^* (p<0.0001); *Col6a2^+/-^*vs *Col6a2^-/-^* (*p<*0.0001). **(G)** Muscle cryosections were stained with fluorescent wheat germ agglutinin (WGA--green) and DAPI (blue) after downhill running and systemic administration of Evan’s blue dye (EBD). Red fibers (EBD positive) represent those with damaged sarcolemma. Scale bar = 50 µm. **(H)** Quantification of EBD positive fibers normalized to muscle section area. For statistical comparisons, ordinary one-way ANOVA was performed at baseline (no differences) and after downhill running with Bonferroni’s adjustment for multiple comparisons. Wildtype (Male n=7, Female n=6); *Col6a2^+/-^* (Male n=4, Female n=5); *Col6a2^-/-^*(Male n=6, Female n=7). **(I)** Tibialis anterior muscle sections stained with H&E after cardiotoxin injury. Regeneration after cardiotoxin injury is delayed in *Col6a2^-/-^* mouse muscle with persistent inflammatory infiltrates (day 7 and day 14), myofiber/myotube smallness, and increased endomysial spaces. Myofibers with internal nuclei represent newly generated myofibers/myotubes after injury. Scale bar = 50 µm. Error bars in all panels represent SEM.

### *Col6a2^-/-^* muscle regeneration after cardiotoxin injury is delayed

Previous studies of other *COL6-*RD mouse models have noted an inadequate muscle regenerative response to injury^23, 24^. To assess the regenerative capacity of *Col6a2^-/-^*mouse muscle, we intramuscularly injected cardiotoxin (CTX) into the mouse tibialis anterior muscle. Skeletal muscle undergoes degeneration after CTX injection, followed by inflammatory cell infiltration, and regeneration of new myofibers marked by internal nuclei. Although regeneration continues until 28 days after injury, the majority of new myofibers are formed by day 14^26, 27^. After CTX injury, *Col6a2^-/-^*and wildtype muscle fibers underwent degeneration, with infiltration of the muscle with inflammatory cells by day three and regeneration by day seven after injury. However, regeneration in *Col6a2^-/-^*muscle was delayed, with persistent inflammatory infiltrates, increased prevalence of small myofibers/myotubes, and increased endomysial space at 14 days after injury (Figure 3I).

### *Col6a2^-/-^* muscle extracellular matrix is fibrotic, has abnormal morphology, and shows impaired passive mechanical properties

Muscle ECM remodeling and fibrosis are pathologic hallmarks of muscular dystrophies, including *COL6-*RD. We found a marked increase in endomysial fibrillar collagen content based on the Sirius red staining of muscle sections starting at 10 weeks of age and through 60 weeks of age in male and female *Col6a2^-/-^* mice (Figure 4A-B). However, *Col6a2^-/-^* animals at five weeks of age -- a timepoint when they show reduced grip strength, muscle atrophy, and other dystrophic histological changes – did not yet show increased fibrosis. We did not detect a sex difference in development or progression of muscle fibrosis. Scanning electron microscopic evaluation of collagen fibers in decellularized lateral gastrocnemius muscles in *Col6a2^-/-^* mice showed an increase in thickness and variability of collagen fibers in the endomysium and perimysium, consistent with increased fibrillar collagen deposition and disorganization of the muscle ECM (Figure 4C).

**Figure 4.**
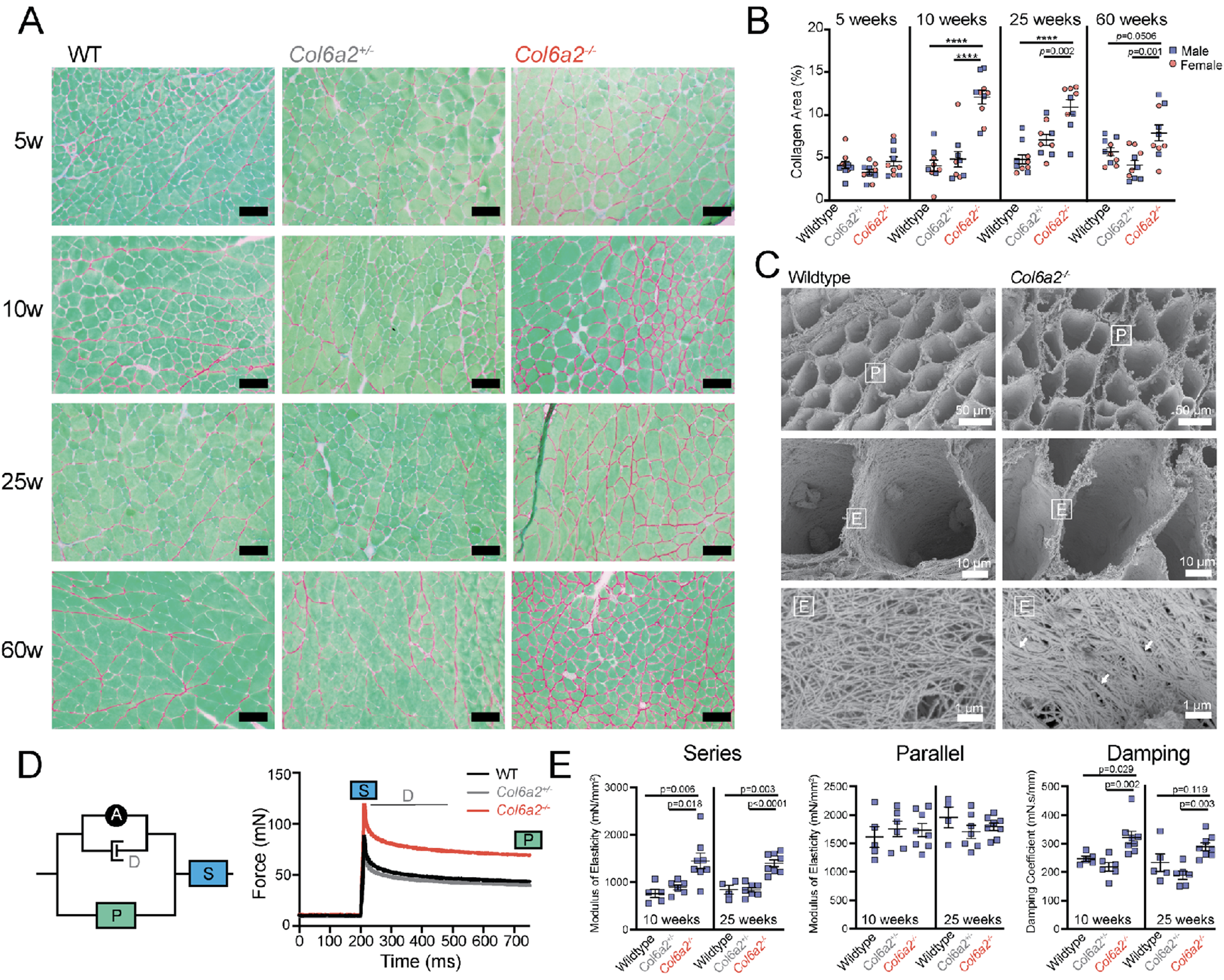
*Col6a2^-/-^*muscle extracellular matrix is fibrotic, has abnormal morphology, and shows altered passive mechanical properties. **(A)** Sirius red stain shows increased fibrillar collagen content (mainly collagen I and collagen III) in *Col6a2^-/-^* mouse muscle, 10 weeks of age or older. **(B)** Quantification of the sirius red positive area (collagen area) in whole muscle sections. **(C)** Scanning electron microscope images of decellularized lateral gastrocnemius muscle of wildtype and *Col6a2^-/-^* mouse shows altered morphology of fibrillar collagens (collagen I). Collagen fibers show increased thickness and variability most notably in the endomysium (arrows). Myonuclear impressions on the endomysium are noted in the top and middle panels. P= perimysium, E=endomysium. **(D)** Schematic of A.V. Hill viscoelastic mechanical model of skeletal muscle. The right panel shows an example of the force tracing of isolated EDL muscle after passive stretch. The different components of the model are calculated from different portions of the stress-relaxation protocol as indicated. S=Series elastic element, P=parallel elastic element, D=damping, A=active contractile element. **(E)** Quantification of series elastic element, parallel elastic element, and damping coefficient in male animals after passive stress-relaxation protocol shows a marked increase in series modulus of elasticity and damping coefficient, but not the parallel elastic element in *Col6a2^-/-^* isolated EDL muscle compared to control littermates. Statistical comparisons were performed by 2-way ANOVA and Tukey’s adjustment for multiple comparisons (panel B) and one-way ANOVA and Tukey’s adjustment for multiple comparisons (panel E). **** = *p<*0.0001. Error bars represent SEM.

We then assessed the functional consequences of fibrosis and ECM remodeling on passive mechanical properties of *Col6a2^-/-^* skeletal muscle. We adapted an experimental model to test viscoelastic properties of EDL muscle based on the AV Hill model^28^. We found a marked increase in the series modulus of elasticity and damping while parallel modulus of elasticity of the model was unchanged (Figure 4D-E).

### TGFβ pathway is upregulated in *Col6a2^-/-^* skeletal muscle

To evaluate the underlying molecular mechanisms of disease in *Col6a2^-/-^*mice, we performed a comprehensive RNA-seq study of muscle tissue in juvenile (five weeks old, four male and four female) and adult (25 weeks old, four male and four female) mice compared to wildtype littermate controls. In the juvenile mice, several ECM and muscle related genes (for example, *Myl4, Lgals3, Sln, Postn, Cilp, Fn*) were upregulated (Figure 5A, and Supplementary appendix 1). In addition, *Mstn*, the gene encoding myostatin, was noted to be downregulated in *Col6a2^-/-^* muscle, a finding consistent with other models of muscular dystrophy^29^. To evaluate the master regulators of the observed transcriptomic signatures, we used the upstream regulator analysis bioinformatic tool (ingenuity pathway analysis [IPA]-Qiagen) which showed a marked upregulation of the transforming growth factor β (TGFβ) pathway and other immune system/cytokine related pathways such as interferon-ɣ and IL-6, among others (Figure 5B). Gene ontology analysis of dysregulated genes using the DAVID database also identified immune system pathways as well as biological processes related to collagen fiber organization and extracellular matrix (Figure 5C). Similar findings were noted in 25- week-old animals, again with the TGFβ pathway as the most upregulated upstream regulator (Figure 5D-F). To scrutinize the apparent upregulation of the TGFβ pathway as a master regulator, we generated heatmaps of expression profile of 70+ TGFβ pathway related genes in each individual sample in the dataset (Figure 5G-H). These results were consistent with a robust dysregulation of TGFβ related genes in *Col6a2^-/-^* muscle, the vast majority of which were upregulated across the different samples including both male and female muscle.

**Figure 5.**
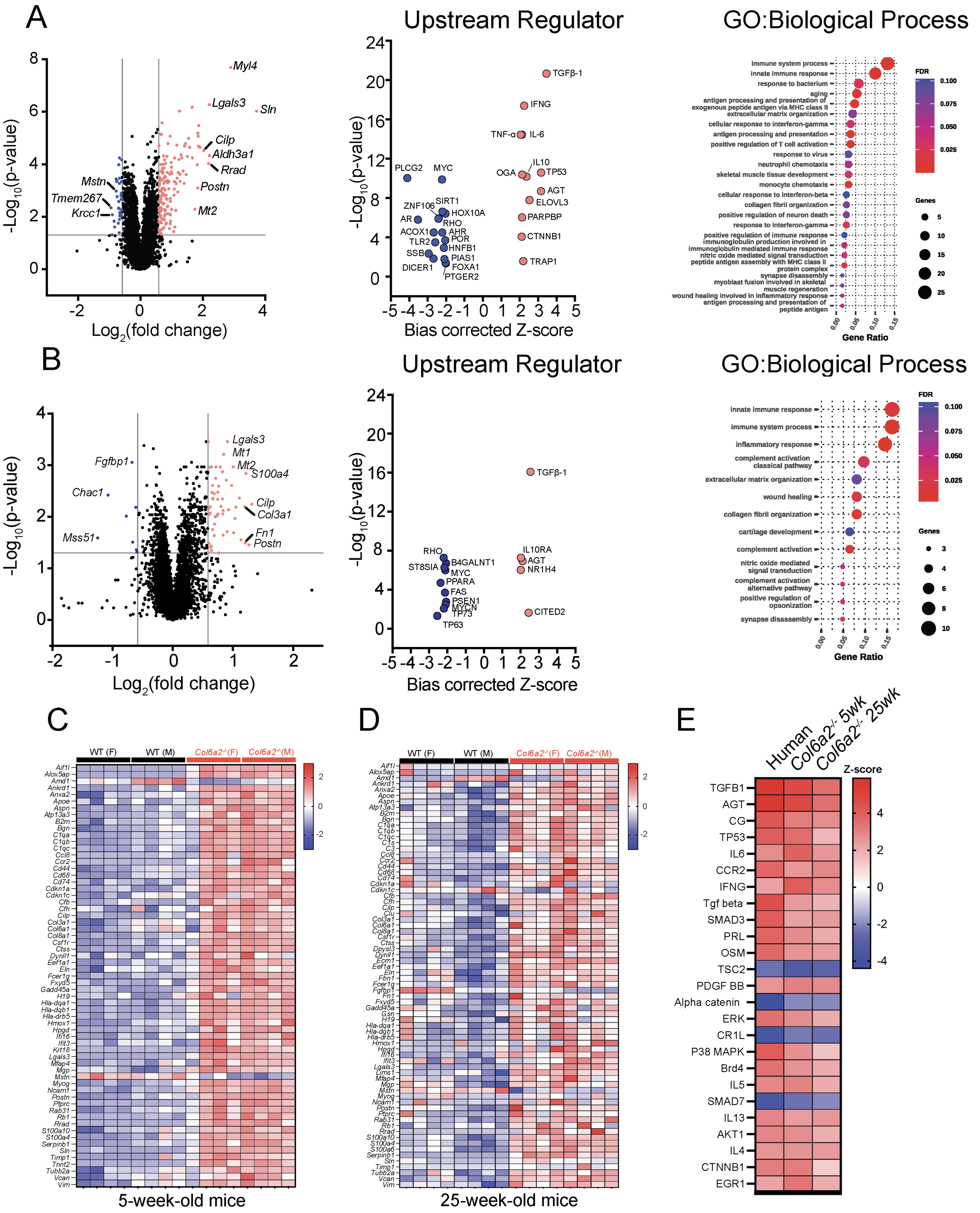
RNA-sequencing identifies a robust upregulation of TGFβ pathway related genes in *Col6a2^-/-^*mouse muscle. **(A)** Volcano plot of differentially regulated transcripts in 5-week-old *Col6a2^-/-^* quadriceps muscle RNA-seq dataset. Upstream regulator analysis of differentially regulated transcripts identified the TGFβ pathway followed by immune system and cytokine related pathways. Gene ontology analysis identified immune system processes and collagen fibril and extracellular matrix related terms. **(B)** RNA-sequencing results from 25-week-old mice shows similar results to A, again with marked upregulation of the TGFβ pathway in the upstream regulator analysis. **(C,D)** Heatmap representation of TGFβ related transcript quantification, (Log_2_ (TPM+2)) depicts the dysregulated transcripts that underlie the TGFβ pathway identification as an upstream regulator. (**E)** Comparison analysis of RNA-sequencing data from the *COL6-*RD human muscle biopsy study (Ref ^30^), 5-week-old and 25-week-old *Col6a2^-/-^* mouse muscle compared to controls. Note upregulation of the TGFB1 pathway in all three datasets.

We have recently reported the upregulation of the TGFβ pathway in muscle biopsies of patients with *COL6-*RD^30^. To compare the transcriptomic profile of muscle tissue in our mouse model with human biopsies, we performed a comparison study of upstream regulators in these datasets using the IPA bioinformatic tools. Upstream regulator and pathways with the most dramatic upregulation or downregulation were common between the human and the *Col6a2^-/-^*mouse datasets (Figure 5I, Supplementary appendix 1).

To validate the transcriptomics findings, we performed western blotting of selected proteins including fibronectin, periostin as well as phosphorylated SMAD3 as downstream intracellular effectors of the TGFβ pathway (Figure 6A, Supplementary Figure 4). These proteins were markedly elevated in *Col6a2^-/-^*muscle lysates. In addition, pSMAD3 expression was notably increased in myonuclei, including internally placed nuclei of regenerating fibers (Figure 6B).

**Figure 6.**
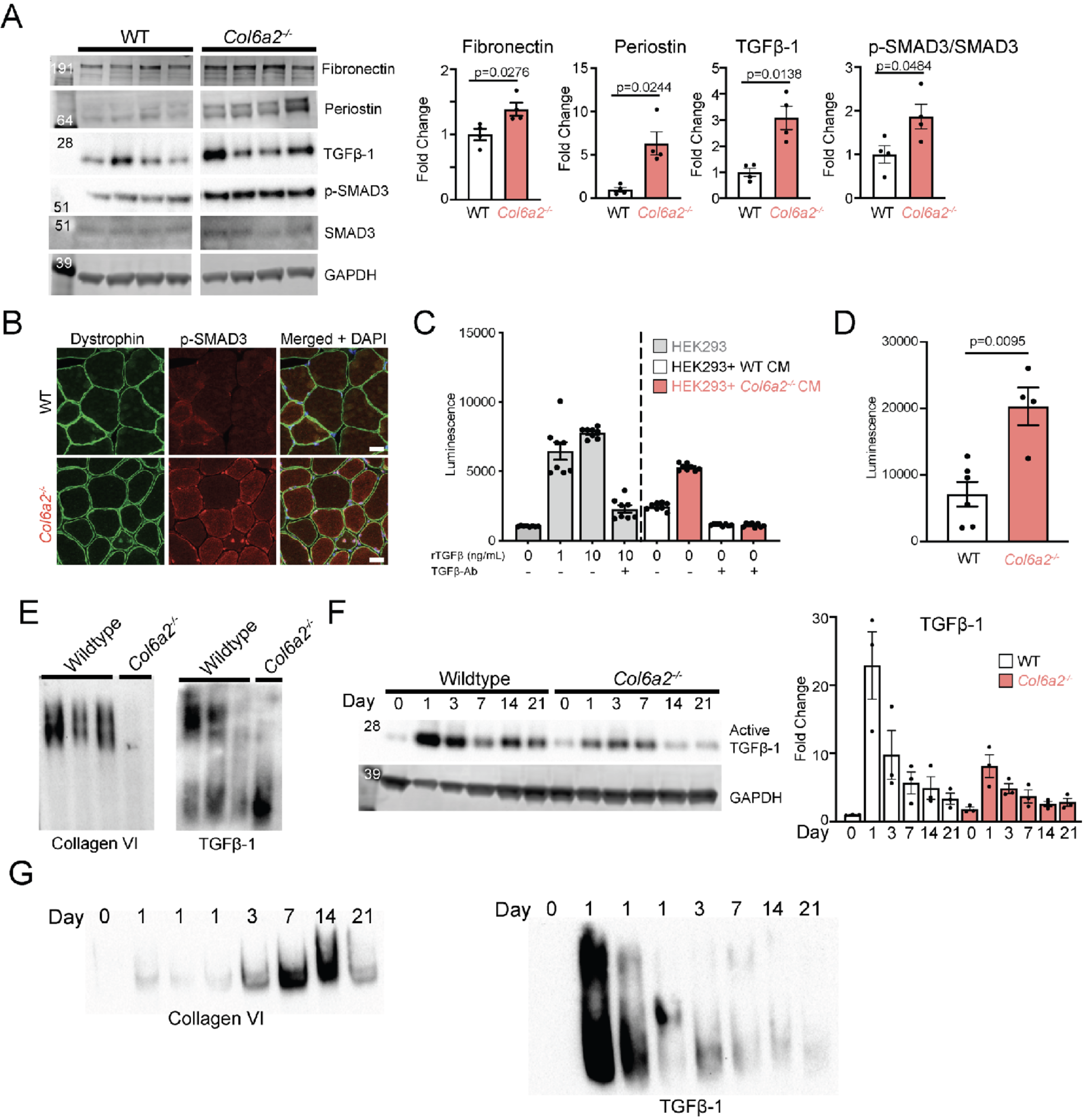
Collagen VI regulates active TGFβ bioavailability. **(A)** Western blotting of mouse muscle lysates under reducing and denaturing conditions show an increase in TGFβ1 and its downstream effectors in soluble fraction of *Col6a2^-/-^*muscle lysates. Corresponding bands were quantified and normalized to GAPDH as an internal control and graphed as fold-change compared to wildtype. Error bars represent SEM. For statistical comparisons a parametric, unpaired *t* test with Welch’s correction was used. **(B)** immunofluorescence staining of muscle tissue shows increased pSMAD3 staining in myonuclei, including internal nuclei of *Col6a2^-/-^* muscle. Scale bar = 25 µm. **(C)** HEK293-luc cells have increased luminescence in response to exogenously added recombinant TGFβ (rTGFβ). This response is diminished with neutralizing antibodies against TGFβ1 (TGFβ-Ab Clone 1D11). Conditioned media from *Col6a2^-/-^* muscle fibroblast cultures results in significantly higher levels of luminescence when added to HEK293-Luc cells compared to wildtype and this response is abrogated by neutralizing antibodies against TGFβ1. The datapoints represent technical replicates (different wells). **(D)** HEK293-luc reporter assay shows that *Col6a2^-/-^*muscle fibroblast cultures deposit higher levels of active TGFβ compared to wildtype controls. Each datapoint is from an independent preparation (biological replicates) and represents the the average of 3-4 technical replicates. **(E)** Gel electrophoresis followed by western blotting of muscle lysates under native conditions (not denatured, not reduced) using anti-collagen VI or anti-TGFβ1 antibodies (as indicated) shows complete absence of collagen VI in *Col6a2^-/-^* mouse muscle. TGFβ1 migrates in a different protein complex in *Col6a2^-/-^* muscle lysates compared to wildtype controls. **(F)** Representative western blots of muscle lysates for TGFβ1 after cardiotoxin injury in soluble muscle lysate fractions. In the wildtype muscle, there is a large increase in active TGFβ1 in the early phases of response to injury (day 1-3), with a subsequent decline in TGFβ levels. In contrast, in *Col6a2^-/-^* muscle lysates, despite elevation of TGFβ1 levels at baseline, they do not increase as much in response to cardiotoxin injury. TGFβ band intensity was normalized to GAPDH as an internal control and compared to the wildtype day 0 (prior to cardiotoxin injury). Data points represent three biological replicates for each timepoint. **(G)** Native gel electrophoresis and western blotting of wildtype muscle lysates after cardiotoxin injury shows an increase in collagen VI expression coinciding with reduced active TGFβ-1 levels, most notable at day 7 and 14 after muscle injury.

### Absence of collagen VI from the ECM increases TGFβ activation

TGFβ isoforms 1, 2, and 3 are three of the canonical TGFβ superfamily ligands that bind and activate TGFβ receptors, which in turn phosphorylate SMAD2/3 that along with SMAD4 translocate to the cell nucleus to induce transcription of the TGFβ pathway-related genes. TGFβ ligands are normally sequestered in the ECM in a latent but poised protein complex and are released in response to specific environmental stimuli^31^. Although the TGFβ pathway was markedly upregulated in *Col6a2^-/-^* mouse muscle, surprisingly, the total transcript levels of three TGFβ isoforms (TGFβ1, TGFβ2, and TGFβ3) were not increased (Supplementary Figure 5). Total TGFβ1 protein levels measured by ELISA in muscle lysates were also unchanged (Supplementary Figure 5). These data suggested that changes in TGFβ activation dynamics instead of its overall transcription and expression underlies TGFβ pathway overactivity in *Col6a2^-/-^* muscle. To investigate the dynamics of TGFβ activation in our mouse model, we first used western blotting and found an increase in the abundance of active TGFβ1 protein in soluble fraction of *Col6a2^-/-^* muscle lysates (Figure 6A). Since latent TGFβ is known to be released by potent detergents, we used 1% NP-40 in these assays in the lysis buffer. In addition, we measured the active TGFβ content deposited in conditioned media of wildtype and *Col6a2^-/-^* muscle fibroblast cultures using a reporter HEK293 cell line that expresses firefly luciferase under the control of a TGFβ/pSMAD2/3 responsive promoter (Supplementary Figure 5C). We first validated the TGFβ responsiveness of the cells (Figure 6C), and then showed a marked increase in TGFβ activity in *Col6a2^-/-^* derived conditioned media compared to controls, which was completely suppressed by addition of neutralizing anti-TGFβ antibodies (Figure 6C,D). Together, these findings suggest that absence of collagen VI from the muscle ECM increases active TGFβ without an increase in total TGFβ transcription or translation.

### Absence of collagen VI alters the TGFβ protein complex

Microfibrillar collagen VI and fibrillin-1 share a similar superstructure^7, 8^. Since fibrillin-1 is known to bind the TGFβ-latent complex in the ECM and thus regulate its bioavailability^32–34^, we hypothesized that similar to fibrillin-1, collagen VI also binds theTGFβ-latent complex. Native gel electrophoresis preserves protein-protein interactions and thus allows for analysis of disrupted protein interactions under different conditions. Using native gel electrophoresis of muscle lysates (not reduced and not denatured) followed by western blotting, we first confirmed the complete absence of the native collagen VI protein in *Col6a2^-/-^* muscle lysates (Figure 6E). We then assessed the TGFβ protein complex in muscle lysates under native conditions and noted a different pattern of migration, with the majority of TGFβ migrating in a lower molecular weight protein complex in *Col6a2^-/-^* muscle lysates compared to controls (Figure 6E, Supplementary Figure 6). This data suggests that absence of collagen VI from the ECM alters the TGFβ complex protein-protein interactions.

### Absence of collagen VI reduces active TGFβ release in response to muscle injury

TGFβ activation from the ECM typically occurs in response to a variety of stimuli such as mechanical force^35–37^ and induced proteolytic activity (for example by plasmin, kallikreins, or matrix metalloproteinases)^31, 38^. Tissue injury, including skeletal muscle injury, causes increased TGFβ activity^39, 40^. Thus, to assess the TGFβ activation and release dynamics from the ECM in *Col6a2^-/-^* muscle, we intramuscularly injected cardiotoxin into the tibialis anterior muscle and compared the TGFβ content in soluble fractions of skeletal muscle lysates exctrated in 1% NP-40 lysis buffer at different timepoints after injury in wildtype and *Col6a2^-/-^* mouse muscle. Wildtype muscle had a large increase in TGFβ levels (>20 fold) within 24 hours of cardiotoxin injection, which gradually declined during the later phases of muscle regeneration (Figure 6F). Although baseline levels of TGFβ were higher in *Col6a2^-/-^* muscle, in the acute phases after cardiotoxin injury, TGFβ release was markedly dampened compared to wildtype controls (Figure 6F, Supplementary Figure 6). In addition, native gel electrophoresis in wildtype muscle lysates after CTX injury showed that native collagen VI levels increase during muscle regeneration, coinciding with decreasing TGFβ levels (Figure 6G, Supplementary Figure 6). These findings suggest that collagen VI regulates TGFβ bioavailability in the skeletal muscle ECM allowing for a robust increase in TGFβ in early phases of regeneration and a gradual decline in later phases of muscle regeneration and myofiber fusion.

### Decreased *Ltbp4-*TGFβ binding does not alter the *Col6a2^-/-^* mouse muscle phenotype

TGFβ is predominantly secreted as part of a large latent complex (LLC) that includes the active TGFβ dimer that is non-covalently bound to its latency associated peptide (LAP) and a latent TGFβ binding protein (LTBP). Four LTBPs are known (LTBP1-4), of which LTBP1, LTBP3, and LTBP4 bind TGFβ with varying efficiencies^31, 41^; however, their absence typically results in reduced TGFβ activity, presumably due to lower efficiency of TGFβ secretion and activation. Ltbp4 is the most abundant Ltbp in mouse skeletal muscle (Supplementary Figure 7) and is a modifier of disease in mouse models of muscular dystrophy^42^ and individuals with Duchenne muscular dystrophy^43^. In addition, stabilization of the hinge region of Ltbp4 protein reduces TGFβ activation and mitigates the muscular dystrophy phenotype in some mouse models of muscular dystrophy^44, 45^.

Since our *Col6a2^-/-^* mice showed altered TGFβ activation in skeletal muscle with evidence of baseline overactivity of TGFβ pathways, we hypothesized that reducing Ltbp4-TGFβ interaction might improve the disease phenotype in our mouse model. We tested this hypothesis by crossing the *Col6a2^-/-^* mice to a knock-in mutant *Ltbp4* mouse model (*Ltbp4^c/s^*) in which a critical cysteine residue in the third TGFβ-binding, 8-cysteine domain of Ltbp4 is mutated^46^. In homozygous mice with this knock-in mutation (*Ltbp4^hom^*), Ltbp4 does not associate with TGFβ-LAP but is normally secreted and localizes to the extracellular matrix^46^. We first re-confirmed the normal localization of Ltbp4 in skeletal muscle in the *Ltbp4^het^* and *Ltbp4^hom^*transgenics and in double homozygote *Ltbp4^hom^/Col6a2^-/-^*mice (Supplementary Figure 7B). We then measured their weight and grip strength, analyzed the muscle histology, and *ex vivo* physiology parameters. Although some of these parameters trended slightly higher, these studies failed to detect a meaningful improvement in any of the functional, physiologic, or histologic assays (Figure 7). This suggests that modulating the Ltbp4-TGFβ interaction is insufficient to address the consequences of TGFβ dysregulation in the *Col6a2^-/-^* muscle ECM.

**Figure 7.**
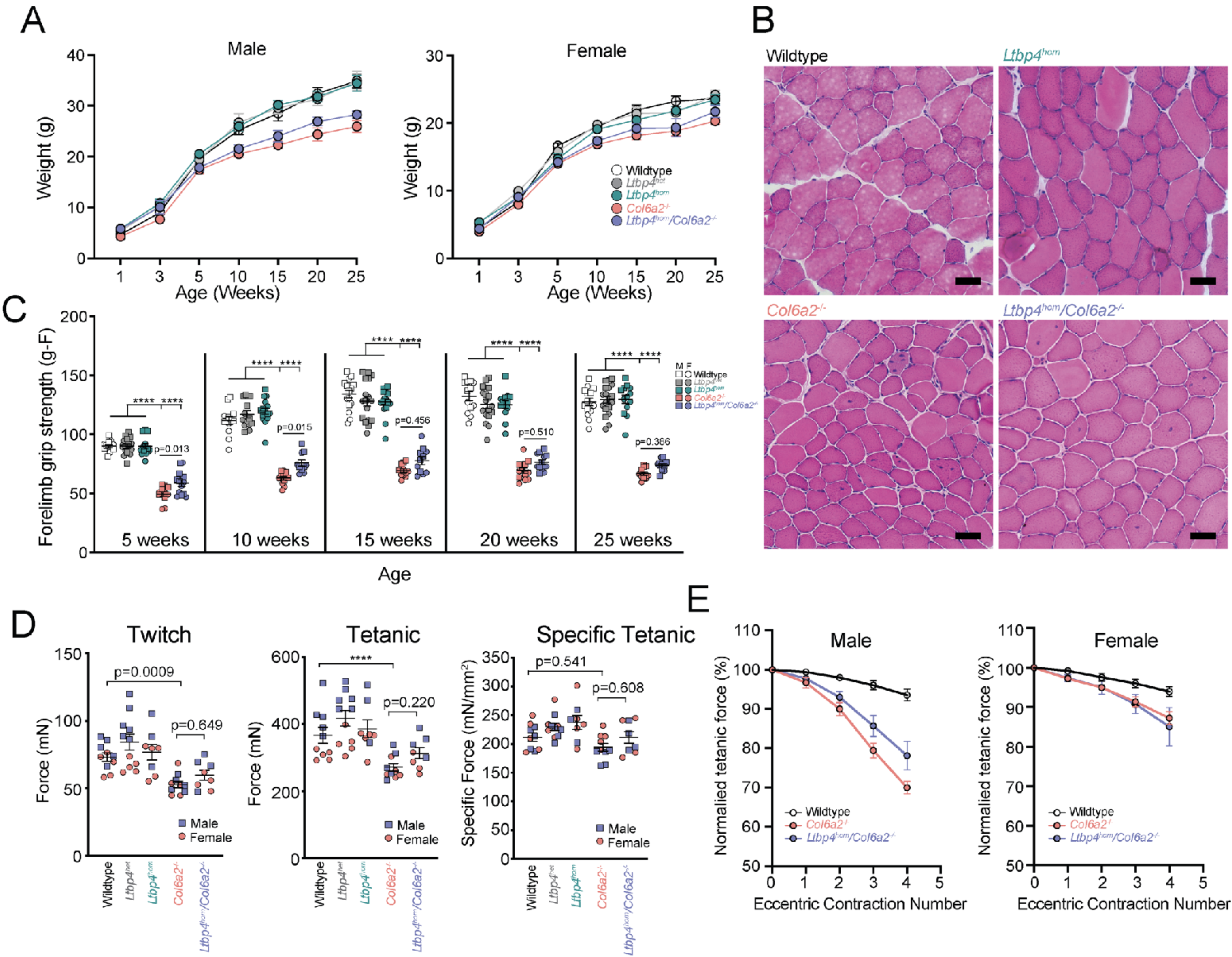
Decreased *Ltbp4-*TGFβ binding does not alter the *Col6a2^-/-^* mouse muscle phenotype. **(A)** Double homozygous *Ltbp4^hom^*/*Col6a2^-/-^* mice weigh similar to *Col6a2^-/-^* mice and are smaller and lighter than wildtype and *Ltbp4^het^* and *Ltbp4^hom^* littermates. **(B)** On hematoxylin and eosin (H&E) staining of tibialis anterior muscle of 25-week-old animals, double homozygous *Ltbp4^hom^/Col6a2^-/-^* muscle tissue shows dystrophic features similar to *Col6a2^-/-^* mice. Scale bar = 50 µm. **(C)** Male and female *Ltbp4^hom^*/*Col6a2^-/-^* mice have slightly higher forelimb grip strength compared to *Col6a2^-/-^* mice, especially in earlier timepoints. However, they maintain markedly reduced forelimb grip strength compared to wildtype and *Ltbp4^het^* and *Ltbp4^hom^*littermates. **(D)** Maximal twitch force and total tetanic force are markedly reduced in 25-week-old *Ltbp4^hom^*/*Col6a2^-/-^* isolated EDL muscle compared to controls and unchanged compared to *Col6a2^-/-^* mouse muscle. The difference in force generation was negligible after normalization to functional cross-sectional area (i.e. specific force). For panels C and D statistical comparisons were performed by 2-way ANOVA and Tukey’s adjustment for multiple comparisons. **** = *p<*0.0001 **(E)** After repeated eccentric contractions, tetanic force declined precipitously in isolated double homozygous *Ltbp4^hom^*/*Col6a2^-/-^* and *Col6a2^-/-^* EDL muscles, more prominently in male animals compared to female animals. Statistical analysis was performed using linear mixed models and Bonferroni adjustment for multiple comparisons. Male: wildtype vs *Col6a2^-/-^* (*p=*0.0008); *Ltbp4^hom^*/*Col6a2^-/-^*vs *Col6a2^-/-^* (*p=*0.469). Female: wildtype vs *Col6a2^--^* (*p=*0.0564); *Ltbp4^hom^*/*Col6a2^-/-^* vs *Col6a2^-/-^* (*p=*1.0). For all panels Error bars represent SEM.

## Discussion

In this work, we report a novel mouse model of *COL6-*RD with homozygous deletion of the *Col6a2* genomic locus resulting in complete absence of collagen VI protein from the skeletal muscle extracellular matrix. Using standardized behavioral tests and physiologic parameters in male and female *Col6a2^-/-^* mice in accordance with Treat- NMD standard operating procedures, when possible, we establish a comprehensive natural history study of this new preclinical model*. Col6a2^-/-^* mice have a normal lifespan but develop muscle atrophy and muscle weakness with histologic hallmarks of muscular dystrophy including myofiber atrophy, rare myofiber degeneration and regeneration, myofiber size variability, and endomysial fibrosis (Figure 1, Supplementary Figure 2). In addition to reduced body weights and isolated muscle weights, we report two standardized, behavioral tests, grip strength and hanging wire test, that show robust differences between control and *Col6a2^-/-^* mice that can be included as readouts in future preclinical and interventional studies.

*COL6-RDs* are a progressive disease in humans; however, we did not note a similar progression in our functional assessments of *Col6a2^-/-^*mice up to 60 weeks of age. Extra-muscular phenotypic studies were not the focus of our current study, but trabecular bone loss has been reported in *Col6a2^-/-^* mice previously^47^ and is a common clinical finding in the human disease. Thus, overall *Col6a2^-/-^* mice manifest with a phenotype that recapitulates the human disease, albeit with less severity and without notable progression.

It is noteworthy that similar to asymptomatic individuals with heterozygous loss of function variants in collagen VI-related genes, heterozygous *Col6a2^+/-^* mice show no signs of muscle weakness or atrophy and perform comparably to wildtype controls in our functional and physiologic studies. In addition, similar to the human disease which does not typically show sex differences in disease severity, in most cases the disease did not affect male or female mice differently; the only exception was relative protection of female *Col6a2^-/-^*mice against eccentric contraction-induced muscle injury compared to males (Figure 3F, Supplementary Figure 3B), a commonly used physiologic parameter in muscle disease research. The underlying mechanism of this finding is not clear and warrants further study, but similar sex differences in this assay attributable to estrogen availability have been reported in female *mdx* mice, a mouse model of dystrophin deficient muscular dystrophy^48^. Thus, our findings confirm that sex differences must be considered in designing future studies that use this physiological parameter as a readout.

Fibrosis is a nearly universal histologic feature in muscular dystrophies. In *Col6a2^-/-^* mice, we noted a gradual increase in endomysial fibrosis over time, first noted in animals 10 weeks of age or older. Ultrastructural studies also demonstrated morphological changes in the endomysial extracellular matrix and fibrillar collagen (predominantly collagen I and collagen III) organization (Figure 4A-C). Importantly, muscle atrophy and weakness at five weeks of age, preceded the appearance of histologically overt fibrosis which was first noted at 10 weeks of age. This suggests that in early phases of the disease the pathogenic processes distinct from muscle fibrosis result in muscle weakness in *Col6a2^-/-^* mice.

We also assessed the passive biomechanical properties of muscle in *Col6a2^-/-^* muscle and found a marked increase in the series modulus of elasticity and damping (Figure 4D-E). The histologic counterpart of these physiologic parameters is not precisely known and includes the intracellular titin protein, the sarcolemma, the myotendinous junction as well as different components of the extracellular matrix^49^; however, the observed perturbations in passive properties of the *Col6a2^-/-^* muscle are most consistent with increased fibrosis and overall collagen content in the ECM. Changes in passive mechanical properties of muscle can contribute to decreased force transmission efficiency^50^. In particular, we noted a very substantial decrease in maximal tetanic force in isolated *Col6a2^-/-^*EDL muscle without a corresponding decrease in specific force (Figure 3C-D), which may be due to decreased efficiency of force transmission through the collagen VI-deficient ECM. Changes in passive mechanical properties of muscle also alters mechanical signal transduction that is important for normal muscle function, and regulation of signaling proteins and growth factors such as TGFβ^51, 52^. Thus, changes in passive mechanical properties of muscle may further contribute to the dysregulation of this and similar pathways that maintain normal muscle function and its regenerative response to injury.

Our unbiased RNA-Seq analysis of mouse muscle tissue identified an upregulation of the TGFβ pathway related genes in *Col6a2^-/-^* muscle both in juvenile (five-week-old) and adult (25-week-old) animals (Figure 5) as the most striking alteration. Upregulation of the TGFβ pathway was also reported as the leading finding in transcriptomic studies of *COL6-*RD human muscle biopsies^30^ and thus provide further validity to our mouse model and its relevance to the molecular mechanisms of the human disease in skeletal muscle.

TGFβ pathway upregulation has been reported in other muscular dystrophies and is thought to drive muscle fibrosis^53–55^. It remains unclear if TGFβ upregulation is an adaptive or maladaptive response to disease-related muscle injury or whether disease-specific mechanisms drive this upregulation. In our study, it is notable that in five-week-old *Col6a2^-/-^* animals, which show clear signs of muscle weakness and atrophy but do not yet show fibrosis at the histologic level, TGFβ pathway was already highly upregulated, suggesting its upregulation precedes fibrosis and thus is associated with early stages of disease pathogenesis in collagen VI deficiency in addition to its pro-fibrotic effects. Consistent with this, in the human transcriptomic study, *COL6-*RD patient muscle biopsies without overt fibrosis also had highly upregulated TGFβ pathway signaling^30^. This is in contrast with similar studies in Duchenne muscular dystrophy that show increased TGFβ pathway signaling in later stages of the disease after an initial inflammatory phase^56^. In addition, overactive TGFβ activity in the extracellular space can contribute to downstream alterations in autophagy flux, mitochondrial abnormalities, and apoptosis ^57^, all of which have been previously reported in *Col6a1^-/-^* mice^21, 22^.

In addition to muscle weakness and fibrosis, *Col6a2^-/-^*mice show increased susceptibility to muscle injury both in the *ex vivo* eccentric contraction paradigm as well as the in vivo downhill running assay (Figure 3F-H, Supplementary Figure 3). Although we only observed rare degenerating myofibers on histological analysis (Supplementary Figure 2), increased internal nuclei in *Col6a2^-/-^*mouse muscle further confirms ongoing myofiber degeneration and regeneration in this model. In addition, similar to *Col6a1^-/-^* mouse models of the disease^23, 24^, *Col6a2^-/-^*mice show decreased regeneration capacity in response to CTX-induced muscle injury (Figure 3I), which likely further contributes to dystrophic features of the disease.

Our data suggests that abnormal bioavailability of TGFβ contributes to the reduced regeneration capacity of *Col6a2^-/-^* muscle. In particular, we noted two distinct alterations in TGFβ bioavailability during muscle regeneration. First, a relative deficiency of active TGFβ in *Col6a2^-/-^*muscle in acute phases of post-cardiotoxin injury (Figure 6F), which coincides with myoblast and fibroblast proliferation and immune cell infiltration of the injured muscle tissue. Relatively reduced TGFβ levels at this juncture may contribute to inadequate myoblast proliferation or alter provisional matrix deposition that is necessary for normal muscle regeneration. Second, a relative increase in active TGFβ in later phases of TGFβ signaling (Figure 6F), which coincides with myotube fusion. Increased TGFβ signaling has been shown to delay muscle regeneration by preventing myotube fusion^39, 58^ and thus may further hamper the normal regenerative response in *Col6a2^-/-^*muscle and is consistent with the delayed regeneration noted in *Col6a2^-/-^*muscle (Figure 3I). Thus, both a relative deficiency as well as a relative overactivity of TGFβ signaling may contribute to the disease pathogenesis in *Col6a2^-/-^* muscle, depending on the physiological demand.

TGFβ activity is predominantly regulated by its release from an ECM-sequestered latent protein complex that is typically associated with microfibrillar components of the ECM. TGFβ activation is locally and temporally regulated and can occur by proteases, mechanical force, or by other proteins or changes in the physiochemical environment^31^. Our data suggests that increased TGFβ activity in *Col6a2^-/-^* muscle is partly due to alteration of TGFβ complex protein interactions (Figure 6E) without significant change in transcription and translation of TGFβ in muscle tissue (Supplementary Figure 5). We thus surmise that the collagen VI deficiency in the ECM interferes with the proper regulation of active TGFβ availability; this in turn results in a relative deficiency of TGFβ when it is in high demand and its overactivity when it is to be sequestered again. However, the exact molecular mechanisms of how absence of collagen VI from the ECM may interfere with the proper regulation of TGFβ complex activation or its sequestration remain unclear.

Based on the growing body of literature linking LTBP4 function and stability to muscular dystrophy phenotype^42–45^, we tested the hypothesis that germline reduction of Ltbp4- TGFβ binding ameliorates the muscular dystrophy phenotype in *Col6a2^-/-^* mice but failed to show notable improvements in muscle strength, histology, or physiological parameters in our mouse model (Figure 7). These negative data suggest that TGFβ overactivity in *Col6a2^-/-^* mouse may occur downstream of Ltbp4-TGFβ binding. Alternatively, other TGFβ-binding Ltbps (Ltbp1 and 3) may compensate for abrogation of Ltbp4-TGFβ binding and thus dampen its potential therapeutic effects. Alternatively, persistent TGFβ availability after its activation and release from the ECM may contribute to TGFβ overactivity. Collagen VI is known to bind two TGFβ binding proteins, biglycan and decorin^59^. Thus, it is possible that collagen VI-biglycan or -decorin complexes work as a post-activation regulator of TGFβ bioavailability and their disruption leads to TGFβ overactivity in *Col6a2^-/-^*mice.

In summary, our study establishes and validates a new mouse model of *COL6-*RD with direct phenotypic and molecular relevance to the human disease and proposes that the collagen VI deficient matrix results in a primary and substantial dysregulation of dynamic TGFβ bioavailability as a specific major contributor to the disease pathogenesis. This study paves the way for future mechanistic studies into the role of ECM in disease pathogenesis in muscular dystrophy as well as interventional studies that target the TGFβ pathway as a therapeutic target in *COL6-*RDs.

## Methods

### Animals, housing conditions, and ethical approval

The mouse strain used for this research project, C57BL/6N-*Col6a2*^tm1(KOMP)Vlcg/MbpMmucd^, RRID:MMRRC_047168-UCD, was obtained from the Mutant Mouse Resource and Research Center (MMRRC) at University of California at Davis, an NIH-funded strain repository, and was donated to the MMRRC by The KOMP Repository, University of California, Davis, originating from Kent Lloyd, UC Davis Mouse Biology Program. The ES cell line was originally donated to the MMRRC by The KOMP Repository, University of California, Davis, originating from David Valenzuela and George Yancopoulos, Regeneron Pharmaceuticals, Inc.

The *Col6a2* ^tm1(KOMP)Vlcg^ mice were backcrossed 5 generations onto a C57BL/6J background, and maintained at the intramural NIH Bldg 35 vivarium. Regular light–dark cycles (12 hour) were maintained. The room temperature was maintained at 20–25°C, with a relative humidity of 40–65%. Untreated drinking water and chow was provided *ad libitum* for the duration of the study. Heterozygous (hereinafter *Col6a2^+/-^)* breeders were used to generate wildtype controls, *Col6a2^+/-^*controls/breeders, and homozygous *Col6a2* ^tm1(KOMP)Vlcg^ (*Col6a2^-/-^*) transgenic animals.

The *Ltbp4^c/s^* knock-in mice were generated by InGenious Targeting Laboratory as previously described^46^ with a substitution of cysteine for serine at aminoacid position 1260 of *Ltbp4* and re-derived at the NIMH transgenic mouse core from mouse sperm straws kindly provided by Dr. Daniel Rifkin’s laboratory at NYU.

Genotyping of the mice was performed using PCR and agarose gel electrophoresis with the primers listed in the table below. The *Ltbp4^c/s^* genotyping was performed by PCR, restriction digest by *BstAPI,* which only digested the wildtype allele, and gel electrophoresis.

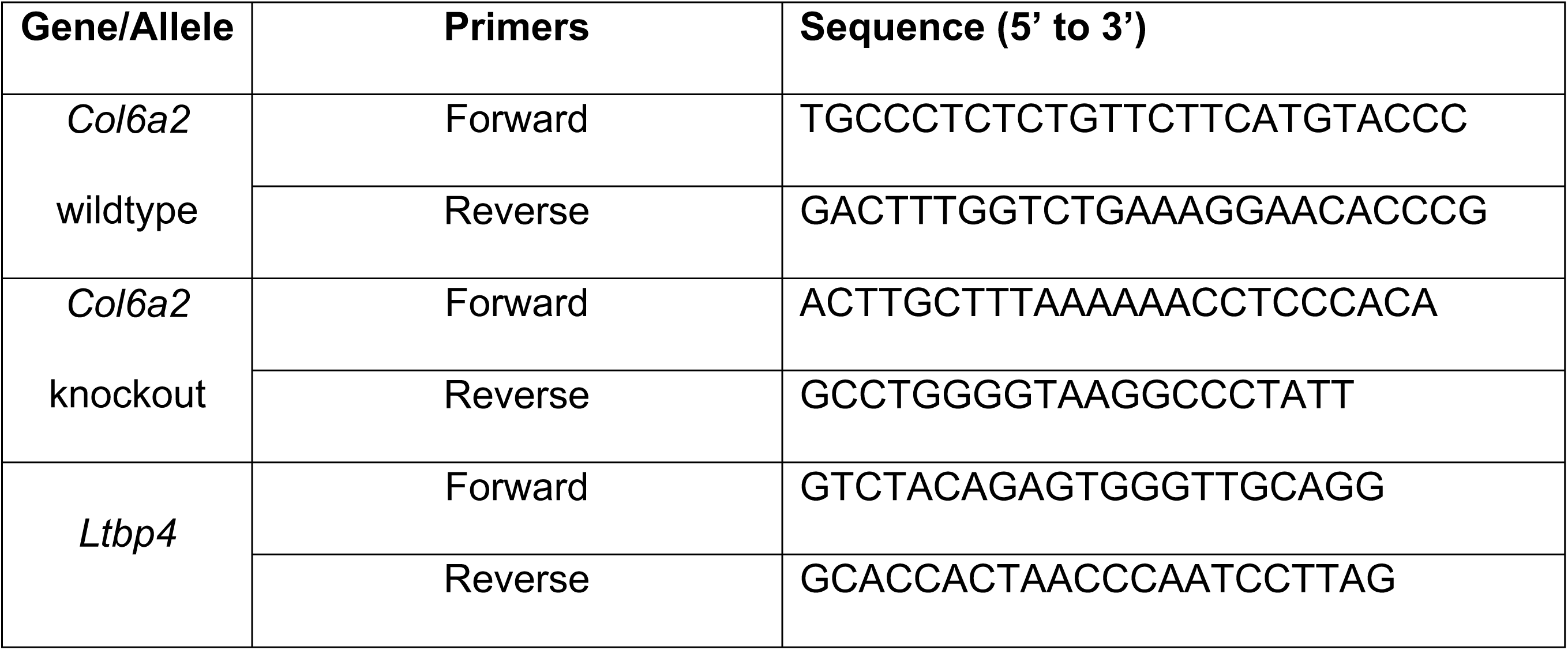

### Functional and *in vivo* studies

Animals were weighed prior to each test. The genotype of the mice was originally masked; however, due to the small size and apparent weakness of the homozygous animals, masking became ineffective. Functional measurements were performed on the same equipment, in the same room, and by the same operator in the same 2-hour window of the light cycle.

#### Grip strength

Forelimb grip strength was assessed using a grip strength meter with a mouse grid attachment (Bio-GS3, Bioseb) and in accordance with TREAT-NMD standard operating procedure MDC1A_M.2.2.001 Ver 1. At each timepoint, six peak force measurements were obtained for each mouse, with at least 1 minute of break in between trials, and averaged.

#### Hanging wire test

Hanging wire test was performed in accordance with TREAT-NMD standard operating procedure DMD_M.2.1.004. Animals were suspended by their forelimbs on a horizontal ∼1.5 mm thick, 55 cm long metallic wire secured by two vertical posts, 35 cm above padded ground and the time when they fell was recorded. The test was stopped if the animal had not fallen after 600 s of suspension. Each animal was tested twice and their best performance was recorded.

#### Rotarod

Rotarod was performed after acclimation of the animals to the equipment (Ugo Basile, Cat# 47600) for a few minutes at low-speed (4-8 RPM) running. For the test, the rotation speed was increased from 2 to 40 RPM over a period of 2 minutes and the time when each animal fell from the apparatus was recorded.

#### Treadmill running, and downhill running paradigm

Treadmill running was performed after acclimation of the animals to the treadmill equipment (Exer 3/6, Columbus Instruments) and running protocol over 2-3 days. For the treadmill running test (at 0°of incline), the animals were run at a speed of 8 m/min for 2 minutes as warmup. The speed was increased by 1 m/min to a maximum of 12 m/min and maintained there until exhaustion. Exhaustion was defined as falling 10 times onto the shocking apparatus. The total distance was recorded. For the downhill running exercise, after the warmup, the mice were run at 15° below horizontal for 30 minutes at 15 m/min. The downhill running was repeated daily over 3 consecutive days. The mice were intraperitoneally injected with Evan’s blue dye (EBD-MP biomedicals, LLC Cat# 151108) at 50 mg/kg body weight prior to running downhill on day 3 and euthanized and dissected 7-8 hours later.

#### Activity meter

A force plate activity monitor (ActiVmeter, Bioseb) with infrared beam detection of rearing was used. The force plate was calibrated before each use and the animal was introduced to the activity cage and monitored for 1 hour, per manufacturer’s specifications. Distance traveled, average speed, and rearing time were recorded every 5 minutes and analyzed using the manufacturer’s software.

#### Voluntary wheel running

Mice were individually housed in cages equipped with a metered running wheel connected to a computer (Scurry Mouse Activity Monitoring wheel, Lafayette Instruments Inc, Model 80820S). Digitized signals were processed by manufacturer’s software (AWM software) and running distance at regular intervals were analyzed. The mice were acclimated to the apparatus for 24 hours and data analysis was performed on day 2 and 3.

### *Ex vivo* force measurements

*Ex vivo* force measurement was performed in accordance with TREAT-NMD standard operating procedure DMD_M.1.2.002. Physiological analysis was performed on isolated EDL muscles using a dual-mode servomotor apparatus and software (Aurora Scientific Inc.) as previously described^60^. Briefly, after dissection, EDL muscle was submerged in a bath of carbogen-equilibrated (95% O_2_–5% CO_2_) Ringer’s solution at 30°C. Optimal length (L_o_) was experimentally determined and measured using calipers. Maximum tetanic force was measured after stimulation of 500 ms duration and of increasing frequency (100-200 Hz). The eccentric contraction stimulation paradigm included a 150- Hz, 700 ms pulse, where a stretch of 10% L_o_ in the last 200 ms of the contraction. Five contractions separated by 3 min of rest in between were administered and the tetanic force prior to the stretch was compared with baseline. The physiologic cross-sectional area was determined as previously described^60^.

Passive mechanical properties of the EDL muscle were measured based on a viscoelastic A.V. Hill model, as previously described^28^. Briefly, after dissection and experimental determination of L_o_, each muscle was subjected to a stretch of 5% L_o_ at a rate of ∼50 L_o_/s, held for 7.0 s before return to L_o_. Series and parallel elastic elements, and damping were then determined using an exponential fit as previously described^28^.

### Histology and light microscopy

Mouse muscles were dissected, frozen in cooled isopentane, and 8 µm cryosections were collected on glass slides and stored at -80 °C. for, Hematoxylin and eosin stain (H&E), sections were incubated in Harris hematoxylin solution (VWR Cat# 95057-858) for 5 minutes, washed with tap water, and incubated with eosin (Sigma Cat#318906) for 1 minute. The slides were then washed in dH_2_O, dehydrated in serially graded ethanol solutions and cleared in xylene, before mounting in Permount™ (Fisher, Cat#SP15-100) and coverslipping.

For the Sirius red stain the sections were incubated in 95% ethanol for 2 minutes, counterstained with 0.5% Fast green (Sigma, Cat# 68724) in PBS (w/v) for 5 minutes, washed with dH_2_O, and incubated with Sirius red solution (0.1% w/v Direct Red 80 in picric acid) for 30 minutes, rinsed with 0.5% v/v acetic acid, and dehydrated serially in graded ethanol solutions, cleared in xylene, mounted in Permount™, and coverslipped. The slides were imaged using a Nikon Eclipse Ti light microscope and tiled images were obtained using a Nikon Digital DS-Fi1 camera.

### Immunofluorescence

Eight µm cryosections were collected on glass slides, fixed in 4% paraformaldehyde for 10-15 minutes at room temperature, and washed in PBS. Sections were blocked with 5% normal goat serum (NGS) or 10% fetal bovine serum (FBS), and 0.5% triton for 1 hour at room temperature before incubation with primary antibodies overnight at 4 °C. After washing with PBS, the sections were incubated with diluted secondary antibodies for 1 hour at room temperature, washed with PBS, and incubated with DAPI solution before mounting with Flouromount-G^®^ and coverslipping. For wheat germ agglutinin (WGA) staining, the sections were incubated in 5 μg/ml Alexa-488 or -633 conjugated WGA in PBS for 10 minutes before mounting. The sections were imaged using a Nikon Eclipse Ti epifluorescence microscope and a Nikon Digital DS-Fi1 camera. Confocal microscopy was performed on a Leica TCS SP5 II confocal microscope. The primary antibodies and dilutions are listed in the table below.

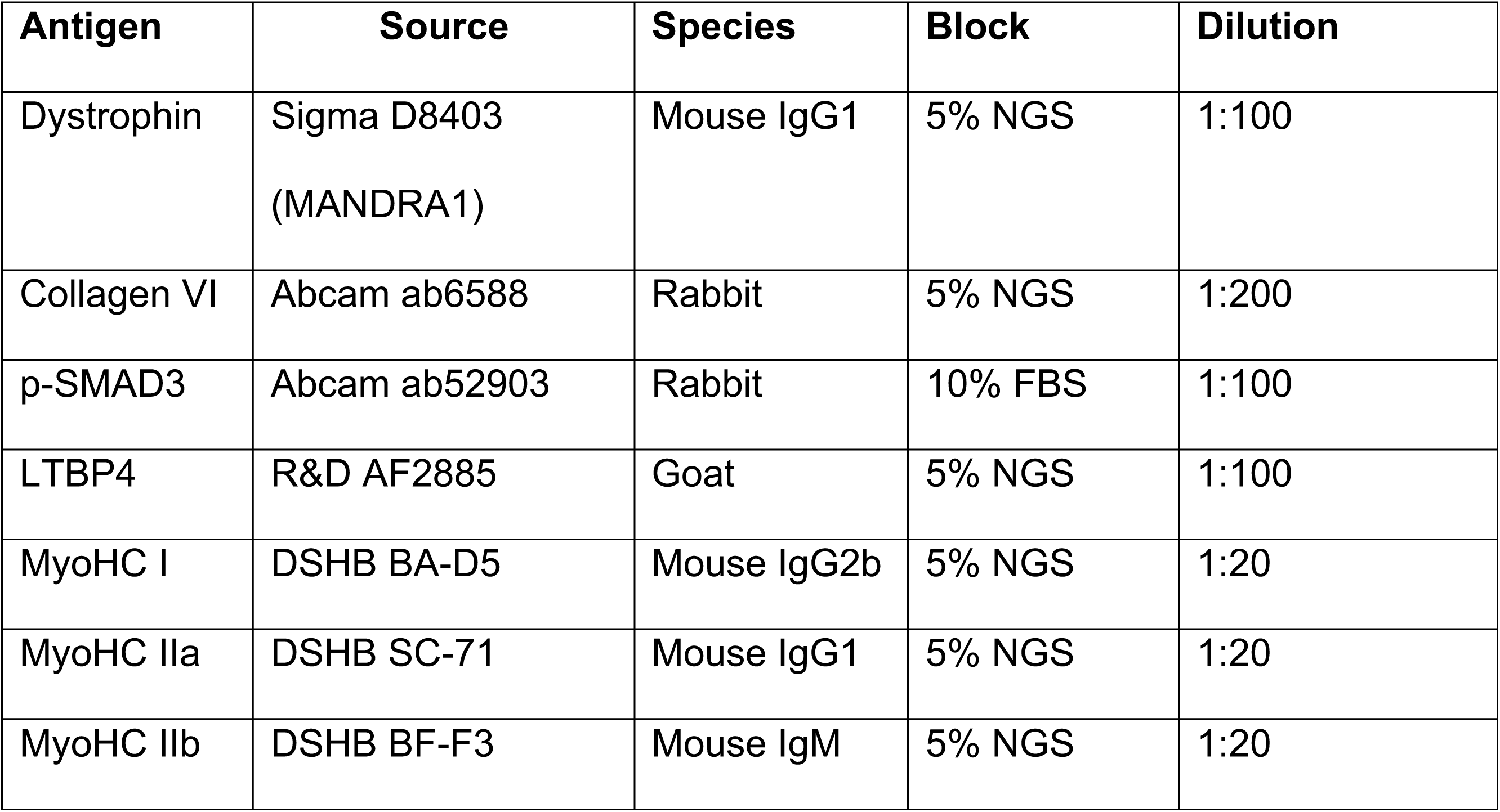

| **Antigen** | **Source** | **Species** | **Block** | **Dilution** |
| --- | --- | --- | --- | --- |
| Dystrophin | Sigma D8403<br>(MANDRA1) | Mouse IgG1 | 5% NGS | 1:100 |
| Collagen VI | Abcam ab6588 | Rabbit | 5% NGS | 1:200 |
| p-SMAD3 | Abcam ab52903 | Rabbit | 10% FBS | 1:100 |
| LTBP4 | R&D AF2885 | Goat | 5% NGS | 1:100 |
| MyoHC I | DSHB BA-D5 | Mouse IgG2b | 5% NGS | 1:20 |
| MyoHC IIa | DSHB SC-71 | Mouse IgG1 | 5% NGS | 1:20 |
| MyoHC IIb | DSHB BF-F3 | Mouse IgM | 5% NGS | 1:20 |

### Quantification of histologic parameters and morphometry

Fibrotic area of the muscle was quantified using the Sirius red stain images by splitting the RGB channels, thresholding the green channel, highlighting the extracellular fibrillar collagen content, and measuring the % area limited to threshold in Fiji/ImageJ. Muscle morphometry was performed using the Myosoft Fiji/ImageJ plugin^25^. Internalized nuclei were manually tallied using the cell counter tool in Fiji/ImageJ.

### Muscle extracellular matrix scanning electron microscopy

The muscle tissue was dissected from euthanized animals, fixed in 4% formaldehyde, and processed as described previously^61, 62^. Briefly, disc-shaped, 2 mm thick pieces of muscle fixed in Karnovsky’s fixative (4% glutaraldehyde, 4% paraformaldehyde, 0.2 mol l−1 sodium cacodylate buffer, pH 7.4) overnight, washed in 0.1 mol/L sodium cacodylate buffer and decellularized in 10% NaOH at room temperature for 4–7 days. The decellularized samples were then prepared for SEM in 1% aqueous tannic acid, and postfixed in 1% aqueous osmium tetroxide overnight. Samples were then dehydrated in ethanol and critical-point dried in liquid CO_2_, mounted on aluminum stubs with carbon tape, sputter coated, and photographed with a Thermo Apreo Volume Scope scanning electron microscope at an accelerating voltage of 2 kV.

### RNA extraction and RNA-Seq

Sixty 10 μm cryosections of quadriceps muscle snap frozen in liquid nitrogen were collected in QIAzole (Qiagen Cat#79306). Four male and four female wildtype and *Col6a2^-/-^* mice were included at two different age groups (5 weeks and 25 weeks old). The RNA was isolated and purified as previously described^63^. RNA quality was analyzed using a 2100 Bioanalyzer (Agilent Technologies, Santa Clara, CA). Samples that had a RIN/RINe ≥ 8 were used for RNA-Seq.

cDNA libraries were prepared according to Illumina protocols TruSeq RNA Sample Prep Kit starting with150 ng of total RNA. The cDNA was fragmented using a Covaris E210 and amplified using 12-15 cycles. Paired-end sequencing of the libraries was performed using the Illumina MiSeq platform with a target depth of 100 million reads of 150-base read pairs per library. Sequence output was processed via RTA and deplexed using CASAVA (http://www.illumina.com/software). Quality inspection of the reads was accomplished using the FastQC tool (https://www.bioinformatics.babraham.ac.uk/projects/fastqc/) and MultiQC tool (https://multiqc.info/) which showed a high quality of sequencing indicated by a GC content peak around 50%, an ambiguity base content of nearly 0%, and an average PHRED score near 40 for all libraries. The Trimmomatic tool (http://www.usadellab.org/cms/?page=trimmomatic) was used (HEADCROP:13, TRAILING:20, SLIDINGWINDOW:4:20, MINLEN:15) to clip Illumina adaptors and trim/remove low quality nucleotides. Intact read pairs were then reference mapped to the Mouse genome (http://daehwankimlab.github.io/hisat2/download/#m-musculus) using HISAT2 (http://daehwankimlab.github.io/hisat2/) under specific conditions (-rna-strandness RF, --fr, --no-discordant, --no-mixed). The returned alignment files were sorted and indexed using samtools (http://www.htslib.org/). Enumeration per gene (Mus_musculus.GRCm38.92.gtf) for each alignment file was accomplished using Stringtie (http://ccb.jhu.edu/software/stringtie/), producing expression in Transcript Per Million (TPM) units. The TPM values were imported into R (http://www.r-project.org/), pedestalled by 2, Log_2_ transformed, and filtered to remove genes not having a transformed value >1 for at least one library/sample. Post-filtering, transformed values for remaining genes were quantile normalized and quality inspected to confirm the absence of outliers via Tukey box plot, covariance-based principal component analysis (PCA) scatter plot, and correlation-based heatmap. Lowess modelling of the normalized data by library class (Coefficient of Variation ∼ Mean) was then performed and the fits were plotted. The lowest mean expression value across the fits at which the linear relationship with Coefficient of Variation was grossly lost (value=2) was defined as the noise threshold for the data. Genes not having a value greater than this threshold for at least one library were discarded as noise-biased. Surviving genes having a value less than the threshold were floored to equal the threshold. Statistical comparisons were performed using Analysis of Covariance (ANCOVA) under AIC-step optimization and Benjamini Hochberg (BH) False Discovery Rate (FDR) Multiple Comparison Correction (MCC) condition, correcting for sex. Genes having a p-value < 0.05 and a linear fold difference of adjusted means > = 1.5X were deemed to be different between sample groups. Volcano plots were generated to describe the number, magnitude, and significance of the genes identified. Gene ontology analysis of the dysregulated genes was performed by using the Database for Annotation, Visualization and Integrated Discovery (DAVID) database (https://david.ncifcrf.gov/). Enriched pathways and functions for the dysregulated genes were identified using the Ingenuity Pathway Analysis tool (https://digitalinsights.qiagen.com/).

The datasets generated and/or analyzed during this study are available in the NCBI repository, https://www.ncbi.nlm.nih.gov/geo. The RNA-Seq data generated as part of this analysis has been deposited in the NCBI Gene Expression Omnibus (GSE228223).

### Muscle Fibroblast Isolation and conditioned media collection

Skeletal muscle fibroblasts were isolated by adapting a previously described protocol^64^. Quadriceps muscle from 3-week-old animals was dissected, minced using sterile razor blades and digested in a 5 mL tube containing 3-5 mL of the digestion mix at 37 °C for 90 minutes: Collagenase type V (Sigma-Aldrich C9263; 5 mg/mL) and Dispase II (Gibco 17105041; 3.5 mg/mL) in DPBS (Gibco 14190094). After the digestion, an equal volume of growth media (IMDM with Glutamax Gibco 31980022, 20% FBS, 1% Chick embryo extract and 1% penicillin/streptomycin) was added to the digestion mix. The neutralized digestion mix was then filtered and spun down to isolate the cells, which were plated in a 10 cm culture dish. After 30 minutes, non-adherent cells were removed, and the adherent cells were used for further studies. Only cells with fewer than 5 passage numbers were used. The adherent fibroblasts were plated at ∼7500 cells/cm^2^ in fibroblast growth media (DMEM/F12, 10% FBS). After 7-10 days, the cells were washed with PBS, F10 media without serum was added (62.5 µl/cm^2^ growth area) and collected after 16 hours, spun down at 300x g for 5 minutes and supernatant (conditioned media) was stored at -80 °C.

### TGFβ activity detection assay

The HEK293 luciferase reporter cell line (HEK293-Luc) was a gift from Dr. Tejvir Khurana (University of Pennsylvania). HEK293 cells were stably transduced with a transgene with a promoter including a SMAD binding element (SBE) (CAGA)_12_ upstream of firefly luciferase transgene. The HEK293-Luc cells were plated at 30,000 cells/cm^2^ in 24-well plates in growth media (DMEM, 10% FBS). After 24 hours, the cells were gently washed with PBS twice. Serum-free F10 medium with or without recombinant TGFβ1 (Peprotech 100-21), conditioned media from wildtype or *Col6a2^-/-^* fibroblast cultures, and/or neutralizing TGFβ antibody (clone 1D11—Invitrogen MA5- 23795) was added. After 16 hours, the cells were lysed with Bright Glo Luciferase assay system (Promega E2610) per the manufacturer’s recommendations and luminescence was measured using a plate reader.

### Protein extraction, gel electrophoresis, and immunoblotting

The muscle tissue was minced and mechanically disrupted using a glass tissue grinder and lysed in approximately 1/10 (w/v), mg of tissue/µl of NP-40 lysis buffer (20mM Tris-HCl pH 7.4, 150 mM NaCl, 100mM EDTA pH 7.4, 1% NP-40; Calbiochem Cat#492016) with protease inhibitors (cOmplete™, Mini, EDTA-free Protease Inhibitor Cocktail Sigma-Aldrich Cat#11836170001) and phosphatase inhibitors (Phostop, Roche Cat# 4906845001). We then centrifuged the extracts at 14,000 g at 4 °C for 15 minutes and collected the supernatant and determined the protein concentration using the BCA protein assay kit (Pierce, Cat#23225). Thirty µg of total protein was reduced and denatured by incubation in LDS sample buffer (Invitrogen NP0007) and 10mM DTT at 95 °C for 5 minutes and separated on a NuPage 4-12% Bis-Tris gel in MOPS buffer and transferred to a PVDF membrane. The membranes were washed, blocked in Intercept blocking buffer (LI-cor Cat#927-70001) or 5% milk in TBS-T (10 mM Tris-HCl pH 7.4, 150 mM NaCl, 0.05% Tween) for 1 hour at room temperature and incubated with the primary antibodies overnight at 4 °C. Bound antibodies were detected with the species-appropriate secondary antibodies by a ChemiDoc XRS (Biorad) imager after enhanced chemiluminescence (Supersignal West Pico Plus, Thermo Fisher Cat#34577) or LI-cor Odyssey DLx scanner for infrared fluorescent tagged secondary antibodies. Densitometry of the bands was performed with Fiji/ImageJ (http://imagej.nih.gov/ij/).

| Antigen | Source | Species | Primary antibody |
| --- | --- | --- | --- |
| Collagen VI | Abcam ab6588 | Rabbit | 1:1000 in 5% BSA |
| p-SMAD3 | Epitomics 1880-1 | Rabbit | 1:500 in 5% BSA |
| SMAD3 | Cell signaling (9513) | Rabbit | 1:500 in 5% BSA |
| TGFβ-1 | R&D systems MAB240 | Mouse | 1:500 in 5% BSA |
| GAPDH | EMD Millipore MAB374 | Mouse | 1:5000 in 5% milk |
| Fibronectin | Sigma F3648 | Rabbit | 1:2500 in Intercept |
| Periostin | Novus NBP1-30042 | Rabbit | 1:1000 in 5% milk |

### Native gel electrophoresis

Protein samples were prepared using the NativePAGE sample preparation kit (Thermo Fisher BN2008) in 1% digitonin or 1% NP-40. Composite 0.5% agarose/ 2.5% polyacrylamide gels were cast as previously described^65^. The gels were run based on NativePAGE system instructions (Invitrogen) with a few modifications. Tris–borate (TBE; 0.45M, pH 8.0) buffer was used in the lower chamber. After adding dark cathode buffer in TBE in the upper chamber, the gel was run at 150 V for 15 minutes on ice. Then, the upper chamber buffer was switched to light cathode buffer in TBE and the gel was run for an additional 50 minutes at 150 V on ice and the proteins were then transferred to a PVDF membrane, which was incubated with 8% acetic acid after transfer for 15 minutes and washed in dH_2_O. The membrane was washed with methanol, before proceeding with immunoblotting.

### Cardiotoxin injury

Mice (∼15 weeks of age) were anesthetized with 2-3% isoflurane. A sterile solution of cardiotoxin (Sigma 217503) in sterile PBS was injected intramuscularly (∼100 μl of 10 µg/ml solution) into the tibialis anterior muscle. The contralateral muscle was injected with ∼100 μl of sterile PBS and used as a negative control and the muscles were collected at different timepoints.

### TGFβ detection by enzyme-linked immunosorbent assay (ELISA)

Total TGFβ1 levels were quantified using the PicoKine TGFβ1 ELISA kit (Boster, cat# EK0515), following the manufacturer’s recommendations. ELISA plates were incubated with sample lysates (5 µg of protein in 100 µl of diluent buffer) or a range of known recombinant TGFβ1 standards in duplicate and incubated at 37 °C for 90 minutes. After discarding the excess mix, biotinylated TGFβ1 antibody was incubated in each well for 60 minutes at 37 °C. Each well was washed three times and incubated with avidin-biotin-peroxidase complex at 37 °C for 30 minutes. Each well was washed 5 times and incubated with color developing reagent for 30 minutes at room temperature, neutralized with stop solution. Absorbance was measured using a plate reader at 450 nm and the values were modeled using a four-parameter logistic curve-fit, to calculate TGFβ1 concentrations.

### Statistics

Two-way analysis of variance (ANOVA) with Tukey’s adjustment for multiple comparisons was used for statistical comparisons, using sex (male or female) and genotype (wildtype, *Col6a2^+/-^*, or *Col6a2^-/-^*) as categorical independent variables. When stratified by sex, ordinary one-way ANOVA with Tukey’s adjustment for multiple comparisons was used for comparisons. Hanging wire test data was analyzed using time-to-event (defined as falling) analysis on a Cox proportional hazard model with Bonferroni-adjusted pairwise comparisons across the genotypes. Eccentric contraction data was analyzed using linear mixed models, with score as the outcome variable and group as the main covariate. Repeated observations were controlled for within subject. Overall group effect was described using F-statistics, and pairwise comparisons between groups. Downhill run data was analyzed using one-way ANOVA. A separate ANOVA was conducted at baseline and after downhill. Western blot protein quantification data was compared using a parametric, unpaired *t* test with Welch’s correction. For the RNA-Seq data, statistical comparisons were performed using Analysis of Covariance (ANCOVA) under AIC-step optimization and Benjamini Hochberg (BH) False Discovery Rate (FDR) Multiple Comparison Correction (MCC) condition, correcting for sex after normalization and noise filtering as described above.

### Approvals

All animal studies were performed in accordance with NIH institutional animal care and use committee approved protocols (#ASP-1337 and #ASP-1469).

## Author contributions

P.M. designed and performed laboratory experiments, analyzed data, and drafted the manuscript. J.R., Y.Z., M.A.N., P.Y., H.H., T.O. and D.A.S. performed laboratory experiments and analyzed data. K.J., G.N., T.O., S.S., T.J.R., D.B.R. analyzed data. C.G.B. designed and oversaw the study, analyzed data, and revised the manuscript. All authors reviewed and edited the manuscript.

## Acknowledgments

Work in C.G. B.’s laboratory is supported by intramural funds from the NIH/NINDS. P.M. and this work was supported by NIH/NINDS career development grant K22NS104135 and a research grant from the University of Pennsylvania Orphan Disease Center in partnership with Cure CMD. D.A.S and T.J.R. are supported by a NIH/NIAMS R01 grant AR055295. The Thermo Apreo VS SEM was purchased with a high-end instrumentation grant from the Office of the Director at the National Institutes of Health (S10OD023461). The authors would like to acknowledge the NIH NHLBI DNA sequencing and Genomics core (Poching Liu PhD, DVM, Yan Luo, PhD) for library preparation and RNA sequencing, the NIH NHGRI microarray core for RNA quality control analysis (Abdel Elkahloun, PhD, and Weiwei Wu), and the NIH NIMH transgenic mouse facility and the NIH Building 35 vivarium staff for providing support for the mouse studies.

---

**Supplementary figure 1.**
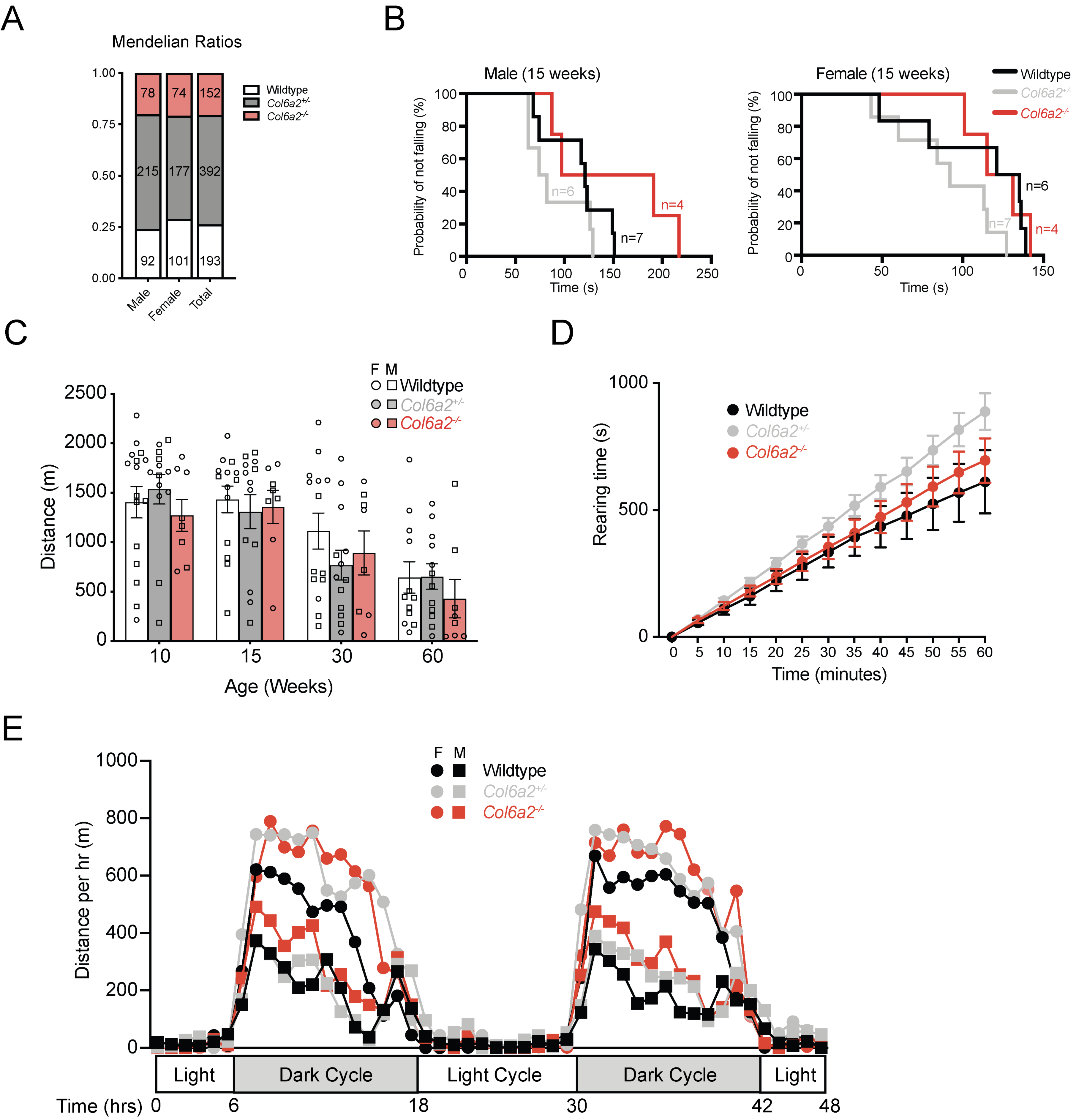

**Supplementary figure 2.**
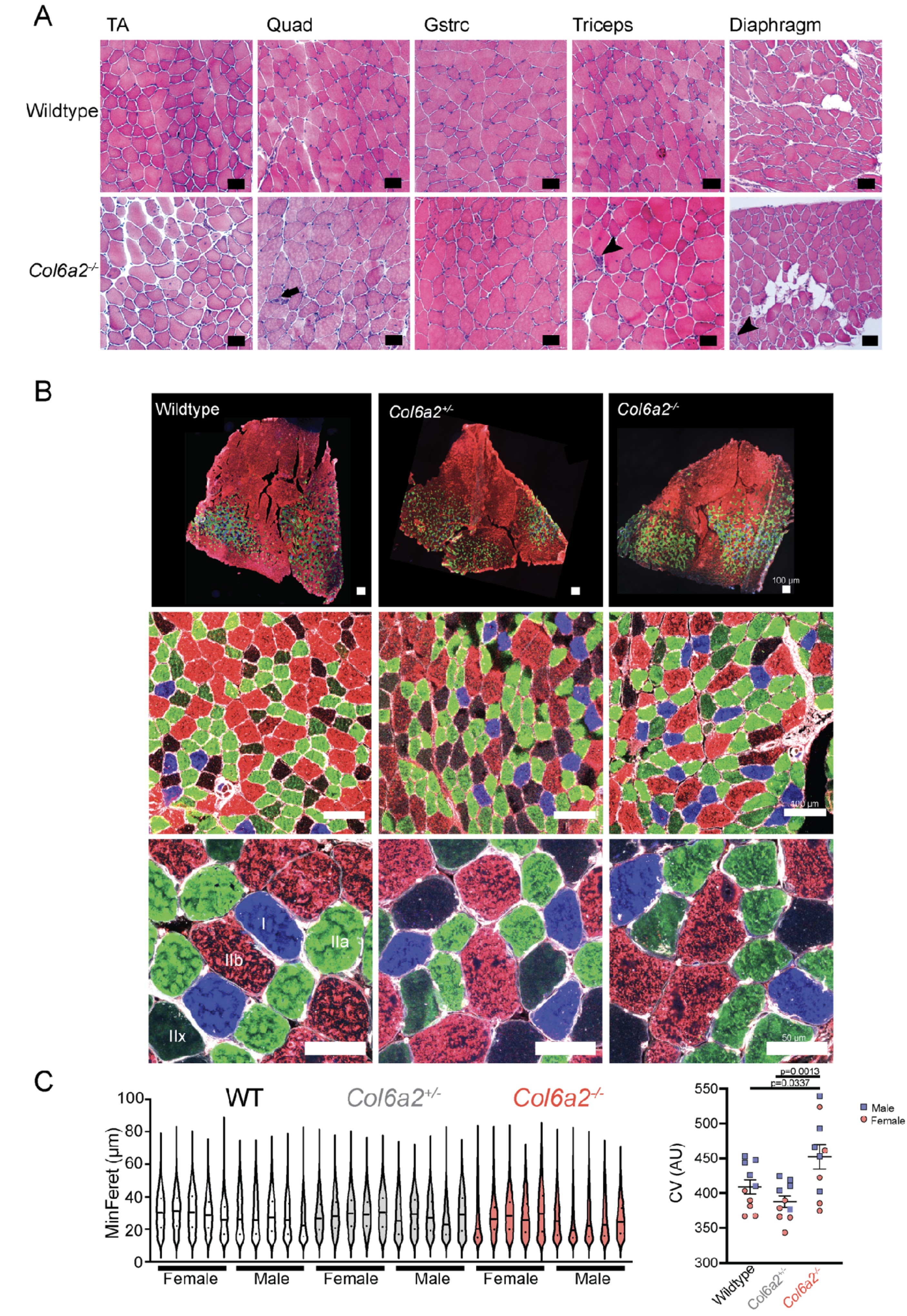

**Supplementary figure 3.**
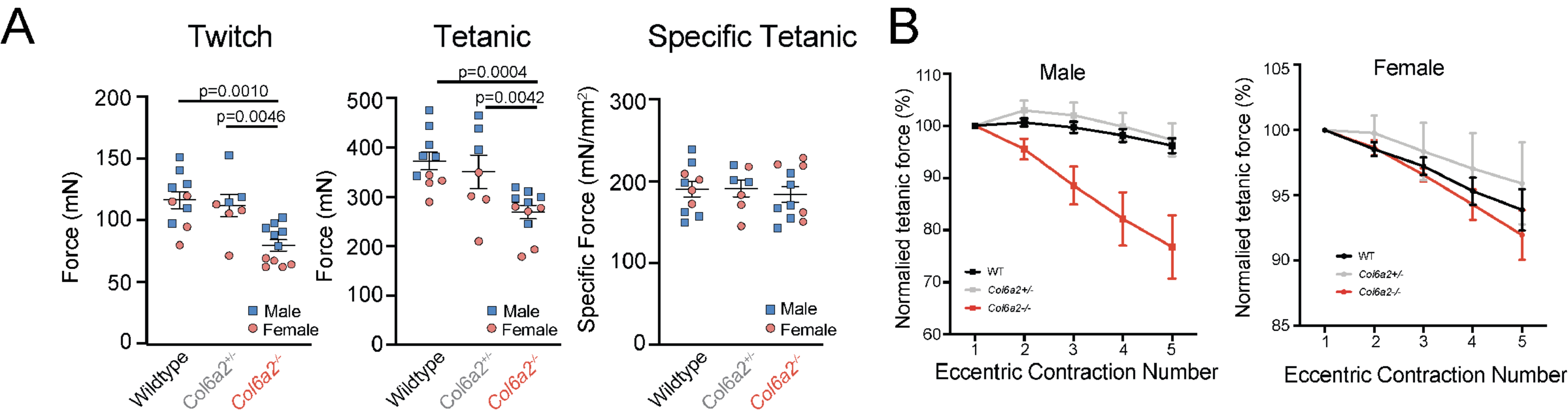

**Supplementary figure 4.**
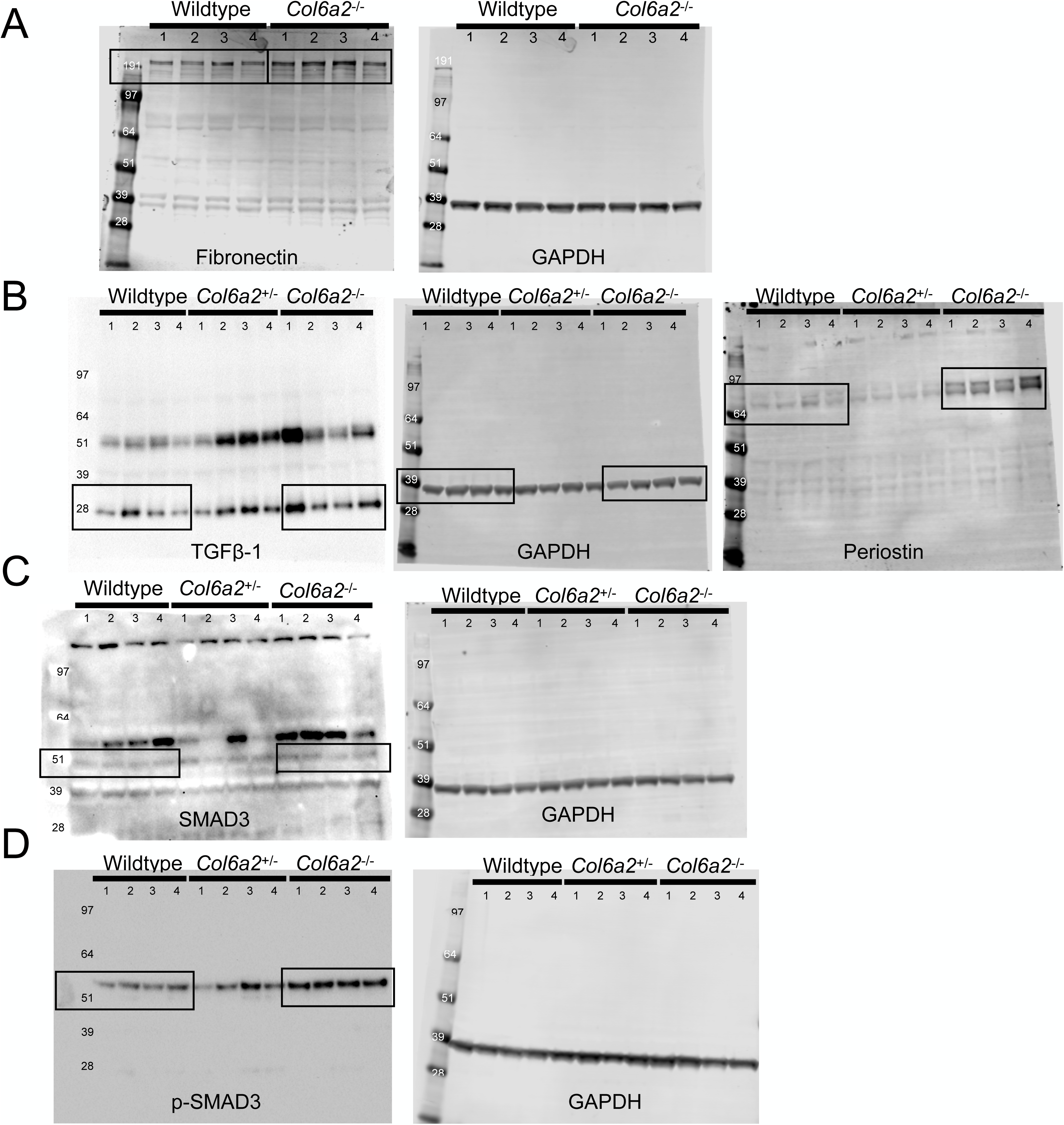

**Supplementary figure 5.**
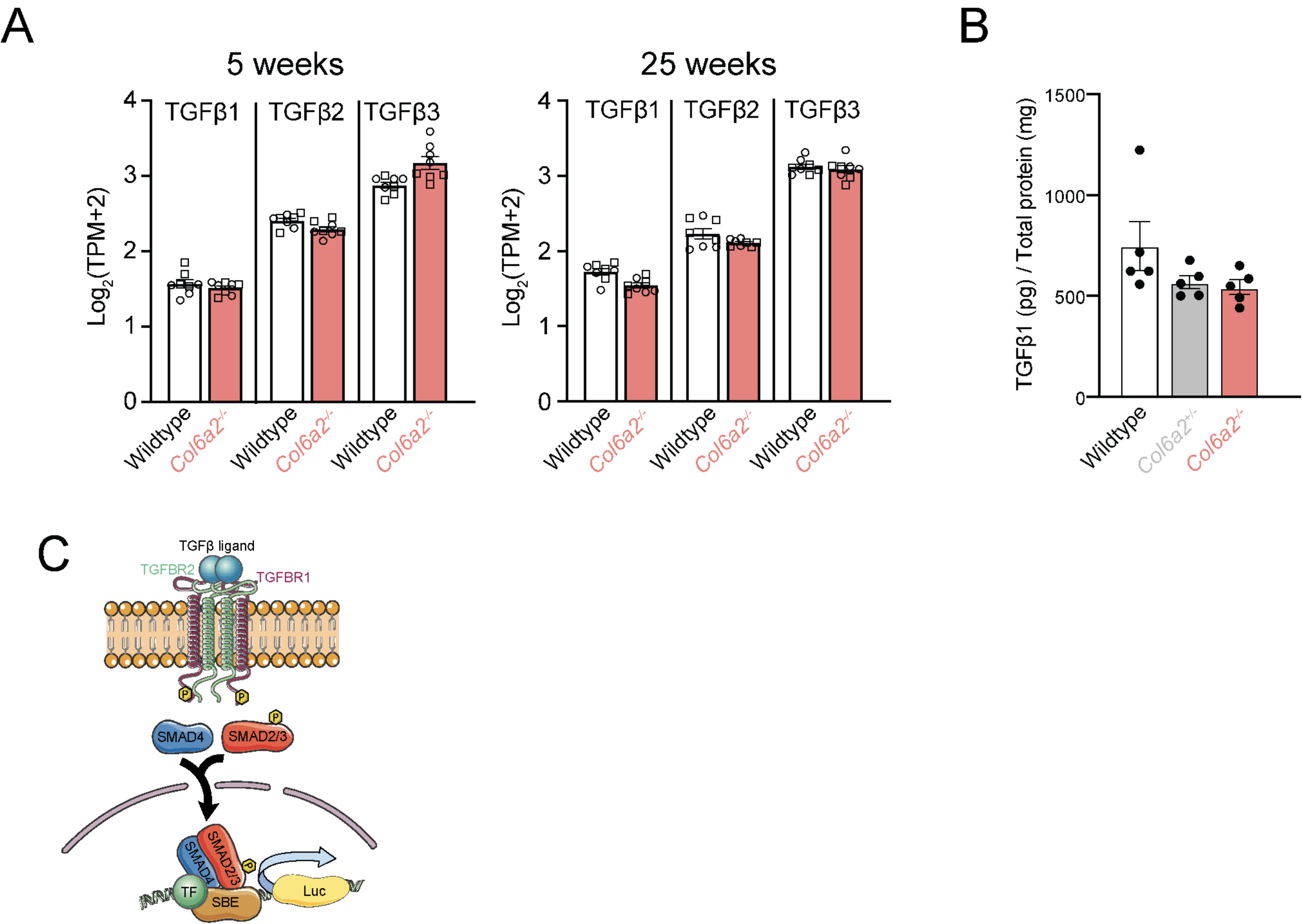

**Supplementary figure 6.**
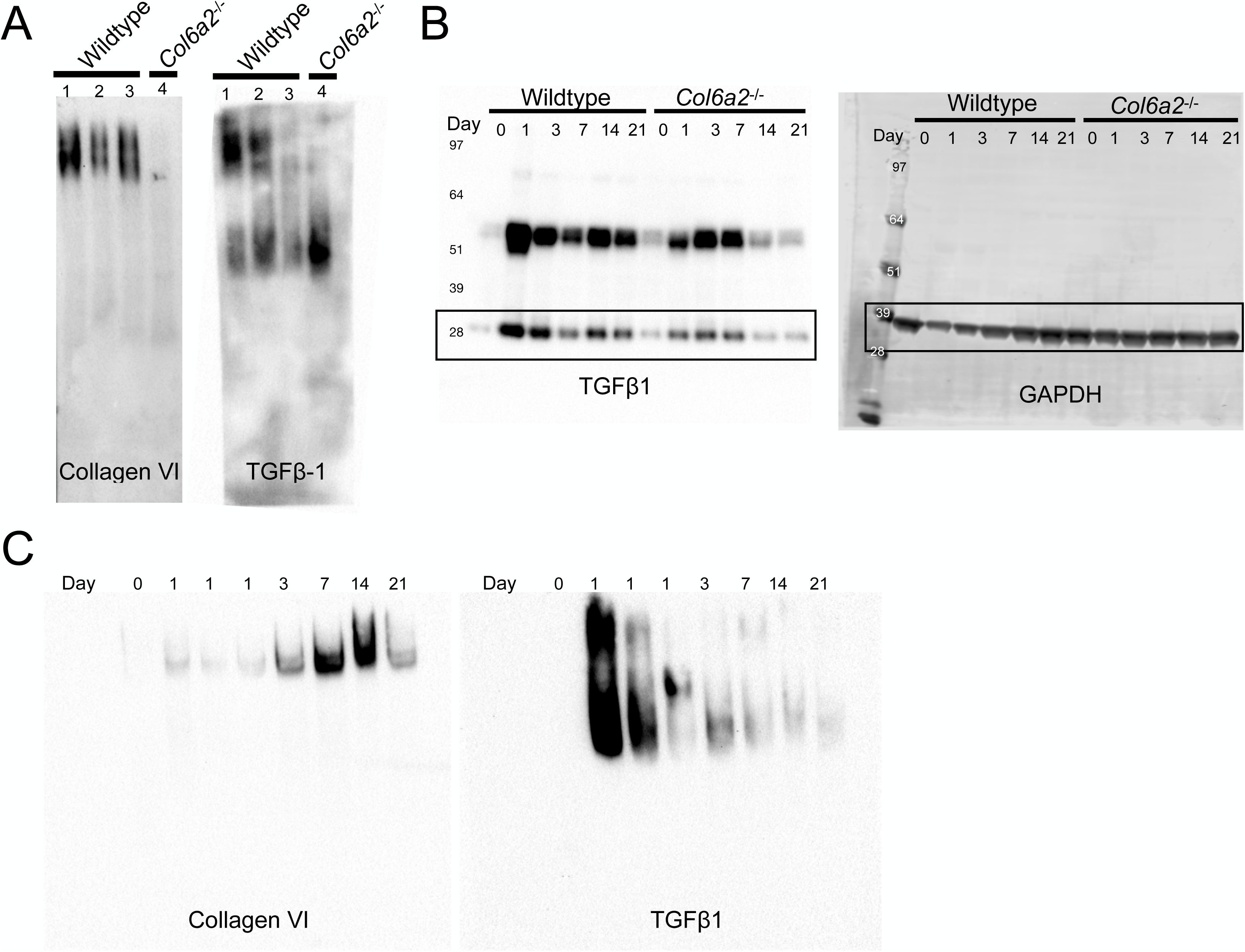

**Supplementary figure 7.**
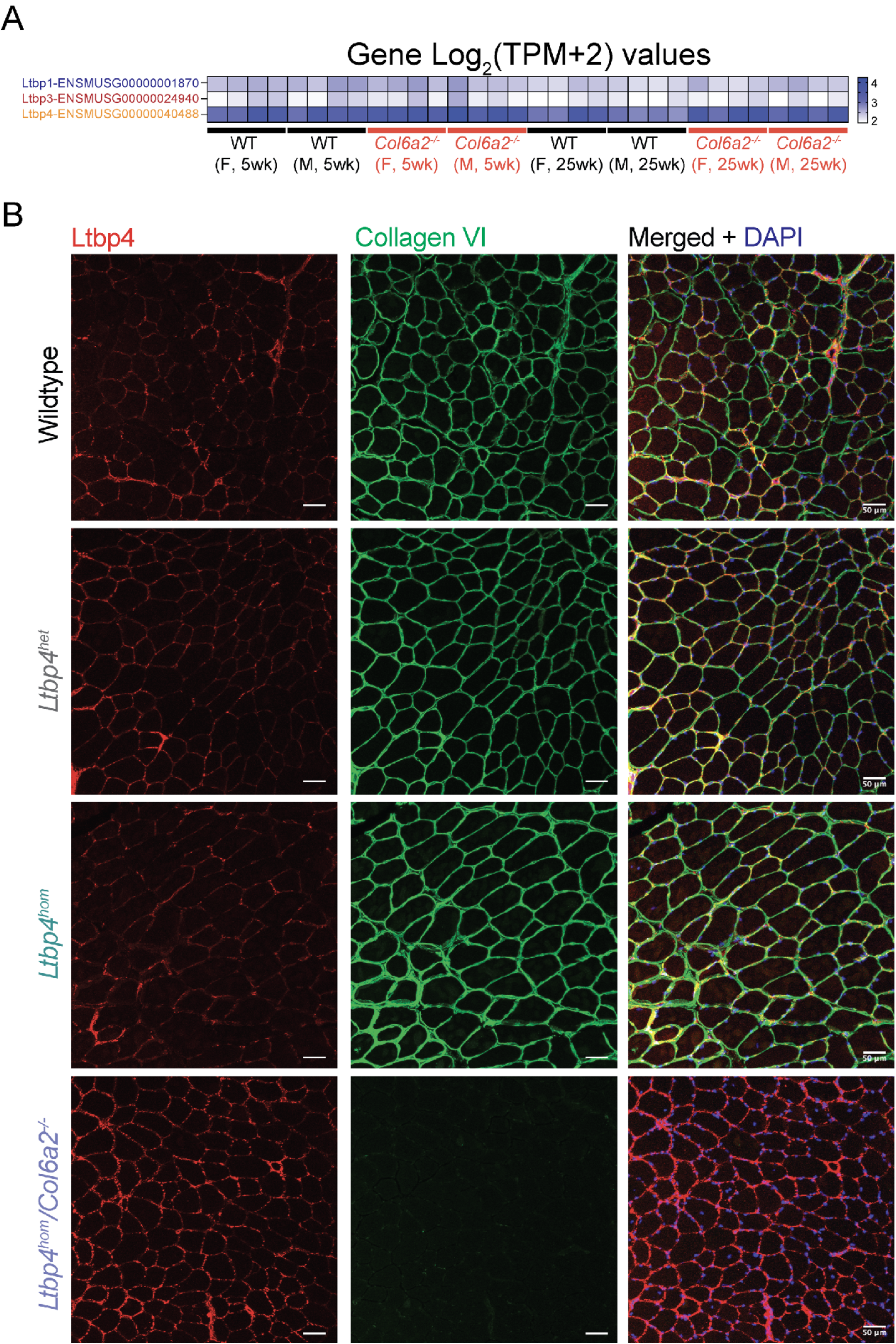

## Notes

**Conflict of Interest:** The authors have declared that no conflict of interest exists.

### Competing Interest Statement

The authors have declared no competing interest.

