## Supplementary Figure for "Collagen type VI regulates TGFβ bioavailability in skeletal muscle"

**Supplementary Figure legends:**

**Supplementary figure 1. (A)** Mendelian ratios from *Col6a2^-/+^* x *Col6a2^-/+^* breeders shows a slight reduction of *Col6a2^-/-^* mice (Male 20.2%, Female 21.0%). **(B)** Rotarod assay fails to show any difference in latency to fall between wildtype, *Col6a2^+/-^* and *Col6a2^-/-^* mice. Similar results were obtained in older mice (up to 60 weeks of age) **(C)** Treadmill running assay fails to show any difference in running distance before exhaustion in wildtype, *Col6a2^+/-^* and *Col6a2^-/-^* mice. Note the large variability among individual mice at each genotype. **(D)** Activity monitoring does not show a difference in rearing time across wildtype, *Col6a2^+/-^* and *Col6a2^-/-^* mice. **(E)** Freewheel running assay fails to show a difference in voluntary running rate across wildtype, *Col6a2^+/-^* and *Col6a2^-/-^* mice. Error bars in panel C and D represent SEM.

**Supplementary figure 2. (A)** hematoxyllin and eosin staining of frozen sections of tibialis anterior (TA), quadriceps (Quad), gastrocnemius (Gastrc), triceps, and diaphragm muscles of wildtype and *Col6a2^‑/-^* mice show dystrophic features with fiber size variability, rare degeneration (arrowhead), and regeneration (arrow) and increased internal nuclei. Scale bar = 50 µm. **(B)** Immunostaining of gastrocnemius muscle for myosin heavy chain subtypes highlighting fiber type I (blue), IIa (green), and IIb (red). Unstained myofibers are considered type IIx. The lower two panels also include wheat germ agglutinin stain (white). There is no qualitative difference or selective loss of specific fiber types in *Col6a2^-/-^* mouse muscle. **(C)** The myofiber minferet diameter of the entire gastrocnemius muscle sections were quantified and shown in violin plots in 60-week-old mice. Note reduced mean fiber diameter in *Col6a2^-/-^* mouse muscle and a marked increase in coefficient of variability (CV) consistent with increased number of atrophic and hypertrophic fibers in *Col6a2^-/-^* mouse muscle. AU= arbitrary units. Error bars indicated SEM. Statistical comparisons were performed by 2-way ANOVA and Tukey’s adjustment for multiple comparisons.

**Supplementary figure 3. (A)** Maximal twitch force and total tetanic force (**C)** are markedly reduced in 60-week-old *Col6a2^-/-^* mouse EDL muscle. The difference in force generation was negligible after normalization to functional cross-sectional area (i.e., specific force). Error bars indicated SEM. Statistical comparisons were performed by 2-way ANOVA and Tukey’s adjustment for multiple comparisons. **(B)** Tetanic force declined precipitously after repeated eccentric contractions in male but not female 60-week-old *Col6a2^-/-^* EDL muscle. Statistical analysis was performed using linear mixed models and Bonferroni adjustment for multiple comparisons. Male: wildtype vs *Col6a2^-/-^* (*p<*0.0001); *Col6a2^+/-^* vs *Col6a2^-/-^* (*p<*0.0001). Female: wildtype vs *Col6a2^-/-^* (p=1.0); *Col6a2^+/-^* vs *Col6a2^-/-^* (*p=*0.216).

**Supplementary figure 4.** Uncropped images of membranes after western blotting for fibronectin (A), TGFβ-1 and periostin (B), SMAD3 (C), and p-SMAD3 (D). Boxes indicate the cropped portions of images in the main figure. Each membrane was also probed with GAPDH as an internal control.

**Supplementary figure 5. (A)** quantification of TGFβ1, 2 and 3 transcripts in mouse muscle tissue from the RNA-sequencing dataset. **(B)** ELISA based quantification of total TGFβ1 levels in muscle lysates from wildtype, *Col6a2^+/-^* and *Col6a2^-/-^* mice shows no marked differences. **(C)** Schematic of the HEK293-luc reporter cell line. In the presence of extracellular TGFβ, TGFβ receptor complex is activated and phosphorylates SMAD2/3 which will subsequently bind SMAD4, translocate to the nucleus, and along with other transcription factors (TF) binds the SMAD-binding element (SBE) promoter and drive luciferase (Luc) transcription.

**Supplementary figure 6.** Uncropped images of Western blot membranes (A) after native gel electrophoresis of muscle lysates probed with antibodies against active TGFβ-1 and collagen VI. Note the complete absence of native collagen VI in *Col6a2^-/-^* muscle. TGFβ1 associated native protein complex in *Col6a2^-/-^* muscle lysates migrates lower on the gel, suggesting disrupted protein-protein interactions. (B) Uncropped images of membranes after western blotting of muscle lysates ad different timepoints after cardiotoxin injury for TGFβ-1 and GAPDH. Boxes indicate the cropped portions of images in the main figure. (C) Uncropped images of Western blot membranes. Protein complexes in wildtype muscle lysates after cardiotoxin injury were separated under native conditions and serially probed with antibodies against active TGFβ-1 and collagen VI. During muscle regeneration, native collagen VI levels increase, which corresponds to decreased levels of active TGFβ-1.

**Supplementary figure 7. (A)** Heatmap representation of transcript leves of three latent TGFβ binding proteins (LTBPs) in the mouse RNA-sequencing dataset in this study. Ltbp1, 3 and 4 bind TGFβ while Ltbp2 does not. Ltbp4 is the most abundant Ltbp in wildtype and *Col6a2^-/-^* mouse muscle tissue. **(B)** Immunoflourescence staining of muscle sections for Ltbp4 and collagen VI. Ltbp4 protein localizes normally to the extracellular matrix in the knock-in *Ltbp4* mouse model or double homozygous *Ltbp4^hom^/Col6a2^-/-^* mice. Scale bar = 50 µm.
